# GPR30 in spinal cholecystokinin-positive neurons modulates neuropathic pain

**DOI:** 10.1101/2024.09.24.614834

**Authors:** Qing Chen, Hui Wu, Shulan Xie, Fangfang Zhu, Fang Xu, Qi Xu, Lihong Sun, Yue Yang, Linghua Xie, Jiaqian Xie, Hua Li, Ange Dai, Wenxin Zhang, Luyang Wang, Cuicui Jiao, Honghai Zhang, Xuelong Zhou, Zhen-Zhong Xu, Xinzhong Chen

**Affiliations:** Department of Anesthesia, Women’s Hospital, Zhejiang University School of Medicine, Hangzhou, 310006, China; NANHU Brain-Computer Interface Institute, Hangzhou, 311100, China; Department of Anesthesiology, The Second Affiliated Hospital, Chongqing Medical University, Chongqing, 400010, China; Department of Anesthesiology, Affiliated Hangzhou First People’s Hospital, Westlake University School of Medicine, Hangzhou, 310006, China; Department of Anesthesiology, Sir Run Run Shaw Hospital, School of Medicine, Zhejiang University, Hangzhou, 317500, China; Department of Anesthesiology, First Affiliated Hospital and School of Brain Science and Brain Medicine, Zhejiang University School of Medicine, Hangzhou, 310058, China; Liangzhu Laboratory, MOE Frontier Science Center for Brain Science and Brain-machine Integration, State Key Laboratory of Brain-machine Intelligence, NHC and CAMS Key Laboratory of Medical Neurobiology, Zhejiang University, Hangzhou, 311121, China

**Keywords:** GPR30, neuropathic pain, spinal CCK positive neurons, corticospinal direct projections, descending facilitation

## Abstract

Neuropathic pain, a major health problem affecting 7-10% of the global population, lacks effective treatment due to its elusive mechanisms. Cholecystokinin-positive (CCK^+^) neurons in the spinal dorsal horn (SDH) are critical for neuropathic pain, yet the underlying molecular mechanisms remain unclear. Here, we show that the membrane estrogen receptor G-protein coupled estrogen receptor (GPER/GPR30) in spinal neurons was significantly upregulated in chronic constriction injury (CCI) mice and that inhibition of GPR30 in CCK^+^ neurons reversed CCI-induced neuropathic pain. Furthermore, GPR30 in spinal CCK^+^ neurons was essential for the enhancement of AMPA-mediated excitatory synaptic transmission in CCI mice. Moreover, GPR30 was expressed in spinal CCK^+^ neurons that received direct projection from the primary sensory cortex (S1-SDH). Chemogenetic inhibition of S1-SDH post-synaptic neurons alleviated CCI-induced neuropathic pain. Conversely, chemogenetic activation of these neurons mimicked neuropathic pain symptoms, which were attenuated by spinal inhibition of GPR30. Finally, we confirmed that GPR30 in S1-SDH post-synaptic neurons was required for CCI-induced neuropathic pain. Taken together, our findings suggest that GPR30 in spinal CCK^+^ neurons and S1-SDH postsynaptic neurons is pivotal for neuropathic pain, thereby representing a promising therapeutic target for neuropathic pain.

## Introduction

Neuropathic pain, a persistent and disabling condition affecting approximately 7–10% of the global population, arises from a lesion or dysfunction of the somatosensory nervous system ^[1–4]^. This disorder manifests as mechanical allodynia (pain triggered by innocuous stimuli) and thermal hyperalgesia (exaggerated pain responses to noxious heat)^[1, 5, 6]^. However, the complex and elusive mechanisms of neuropathic pain still make it difficult to treat. Therefore, identifying precise mechanisms and discovering new therapeutic targets is urgently needed.

The spinal cord (SC) serves as a pivotal hub for processing peripheral sensory inputs and integrating descending modulatory signals. Neuropathic mechanical allodynia occurs via circuit-based transformation in this region^[4, 5, 7, 8]^. Superficial laminae of the SC primarily encode noxious stimuli, whereas deeper laminae process innocuous signals. Under neuropathic conditions, however, low-threshold mechanosensory inputs aberrantly activate nociceptive neurons in superficial laminae through disinhibition and sensitization mechanisms, driving mechanical allodynia. Excitatory interneurons, constituting ∼75% of spinal dorsal horn (SDH) neurons, are central to this pathological process^[9–11]^. Among these, cholecystokinin-expressing (CCK^+^) interneurons, enriched in SDH deep laminae, have recently been proposed as mediators of both mechanical and thermal hypersensitivity^[10, 12]^. Notably, CCK^+^ neurons receive direct corticospinal projections from the primary sensory cortex (S1), modulating neuropathic pain sensitivity^[13]^; however, the molecular underpinnings of this regulation remain obscure.

Estrogen, beyond its classical nuclear receptors (ERα/ERβ), exerts non-genomic effects via the membrane receptor G protein-coupled estrogen receptor (GPR30), which is increasingly recognized as a key player in nociceptive modulation^[14–26]^. Although GPR30’s role in pain modulation has been documented, its specific contribution to SDH circuitry in neuropathic pain remains unexplored.

In this study, we demonstrate that spinal GPR30 orchestrates neuropathic pain by modulating excitability of CCK^+^ neurons and corticospinal descending facilitation. We discovered that GPR30 activation in spinal CCK^+^ neurons was both required and sufficient for the development of neuropathic pain. Interestingly, we observed that GPR30 in spinal CCK^+^ neurons was required for the enhancement of spontaneous excitatory post-synaptic currents (sEPSC) in CCI mice, in an AMPA-dependent manner. Importantly, we revealed that GPR30 was expressed in the spinal CCK^+^ neurons receiving direct projections from S1 sensory cortex, and that GPR30 in S1-SDH post-synaptic neurons was critical for CCI-induced neuropathic pain. These findings establish GPR30 in spinal CCK^+^ neurons as a compelling therapeutic target for neuropathic pain management.

## Results

### Spinal inhibition of GPR30 reverses CCI-induced neuropathic pain and neuronal activation

To investigate the functional relevance of spinal GPR30 in neuropathic pain, we employed a chronic constriction injury (CCI) model to evaluate whether pharmacological blockade of spinal GPR30 alleviates neuropathic pain (Fig 1A). Quantitative PCR (qPCR) analysis revealed a significant elevation of *Gper1* (GPR30) mRNA levels in the lumbar spinal dorsal horn (SDH) of CCI mice compared to sham (Fig 1B). In addition, the *Gper1* mRNA levels in the DRG remained unchanged after CCI (Fig 1B). Intrathecal application of the GPR30 antagonist, G-15, effectively attenuated CCI-induced mechanical allodynia and thermal hyperalgesia in both sexes of mice (Fig 1C-E), whereas basal nociceptive thresholds remained unaltered in naïve mice (Fig S1A and B). Of note, the analgesic effects of G-15 lasted for up to 6 hours after intrathecal application in CCI mice (Fig S1C). Immunochemical data demonstrated that innocuous tactile stimulation triggered pronounced c-Fos expression, a marker of neuronal activation, predominantly in GPR30^+^ cells within the SDH of CCI mice (Fig 1F-J). Critically, G-15 treatment substantially suppressed this c-Fos expression (Fig 1F-J), indicating that spinal GPR30 inhibition normalizes both hypersensitivity and neuronal activation in neuropathic pain in both sexes.

**Figure 1.**
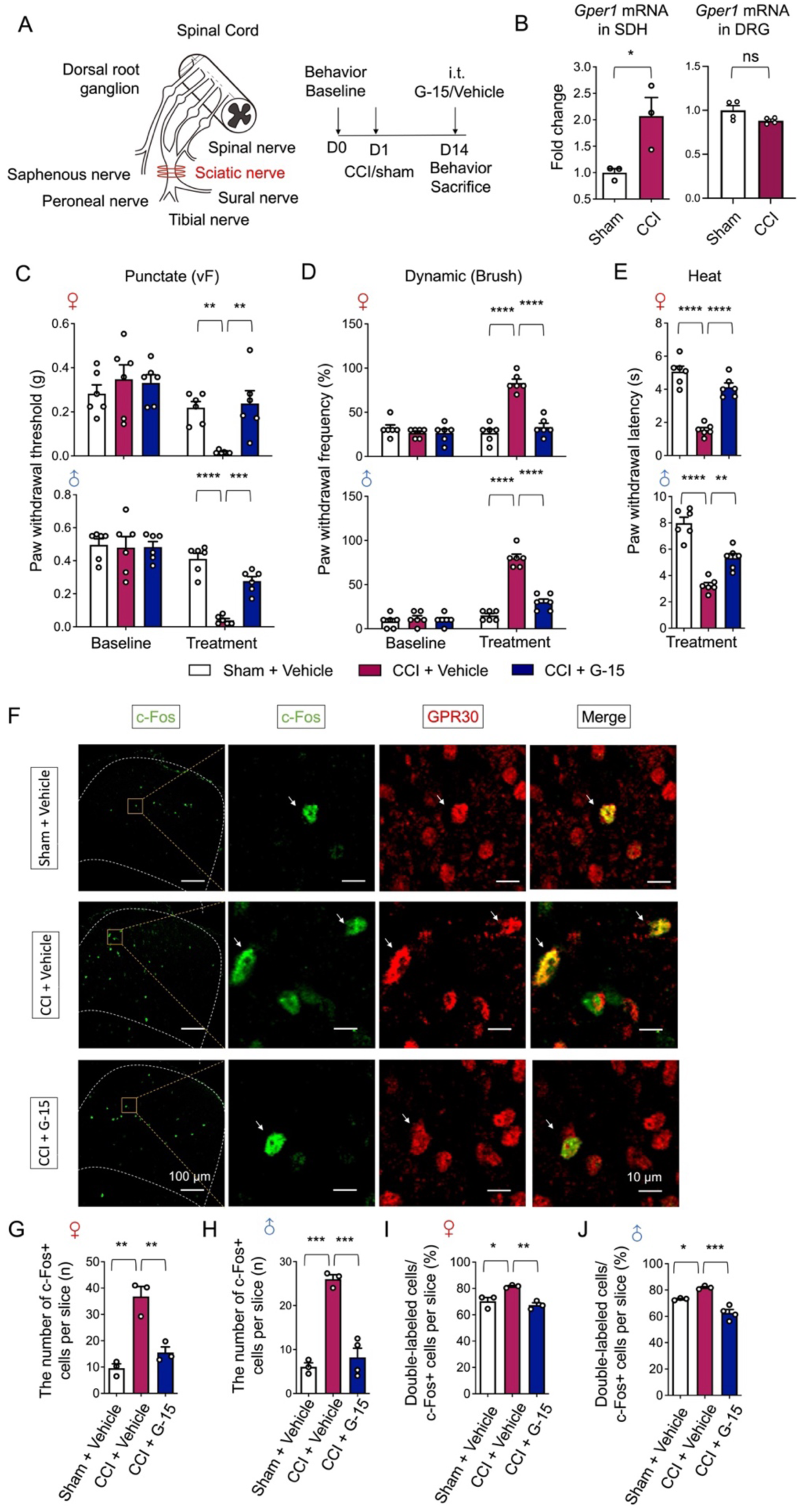
CCI-induced neuropathic pain and neuronal activation were reversed by spinal inhibition of GPR30. **A** Schematic illustration of CCI surgery for induction of neuropathic pain (Left) and diagram showing the timeline of CCI surgery, drug administration and behavioral tests (Right). **B** Quantitative PCR analysis of *Gper1* mRNA in SDH from sham and CCI mice (Left, n = 3 mice for each group) and *Gper1* mRNA in DRG from sham and CCI mice (Right, n = 4 mice for each group). **C-E** Behavioral tests of basic nociception and 14 days after CCI or sham surgery along with intrathecal injection of antagonist of GPR30 or vehicle in Von Frey tests **(C)**, Brush tests **(D)** and Heat tests **(E)** in mice of both sexes (n = 6 mice for each group). **F** Immunochemical detection of c-Fos (Green) and GPR30 (Red). Scale bars: 100 μm. Boxed area of images is enlarged on the right. Scale bars: 10 μm. White arrows indicate double-positive cells. **G, H** Total number of c-Fos positive neurons in the SDH per section in female mice **(G)** and male mice **(H)** (n = 3-4 mice for each group, 4-6 pictures were analyzed for each mouse). **I, J** Percentage of c-Fos positive neurons expressing GPR30 in female mice **(I)** and male mice **(J)**. (n = 3-4 mice for each group, 4-6 pictures were analyzed for each mouse). Data information: in **(B)**, *P < 0.05 (Unpaired Student’s t-test). In **(C, D)**, **P < 0.01; ***P < 0.001; ****P < 0.0001 (two-way ANOVA with Turkey’s multiple comparisons test). In **(E)**, **P < 0.01; ****P < 0.0001 (one-way ANOVA with Turkey’s multiple comparisons test). In **(G-J)**, *P < 0.05; **P < 0.01; ***P < 0.001 (one-way ANOVA with Turkey’s multiple comparisons test). All data are presented as mean ± SEM.

We further probed whether GPR30 activation alone could mimic neuropathic symptoms in naïve mice. Intrathecal application of the selective GPR30 agonist, G-1, robustly evoked mechanical allodynia and thermal hypersensitivity, with symptom persistence exceeding 48 hours (Fig S1D-G). Consistent with behavioral outcomes, G-1 administration markedly increased c-Fos^+^ neuron density in the SDH, with over 80% of activated cells co-expressing GPR30.

Collectively, these findings establish spinal GPR30 as a critical mediator of sex-independent neuropathic pain development in the CCI model.

### GPR30 in spinal CCK^+^ neurons is required for CCI-induced neuropathic pain

To delineate the spinal cellular substrates of GPR30-mediated pain modulation, we first mapped its expression pattern. Our results revealed exclusive GPR30 localization to neuronal populations within the SDH instead of astrocytes or microglial cells via immunochemistry (Fig. S2). Notably, as an estrogen receptor, GPR30 expression showed no sexual dimorphism (Fig S2D). To assess the relative expression of GPR30 in excitatory versus inhibitory neurons, we performed *Ish* and immunostaining with *VGAT*, *Vglut2* and GPR30 (Fig 2A). GPR30 was predominantly expressed on the excitatory neurons of SDH (Fig 2B). Further characterization in wild-type mice using AAV2/9-Camk2-mCherry labeling also confirmed predominant GPR30 enrichment in SDH excitatory interneurons (Fig S3).

**Figure 2.**
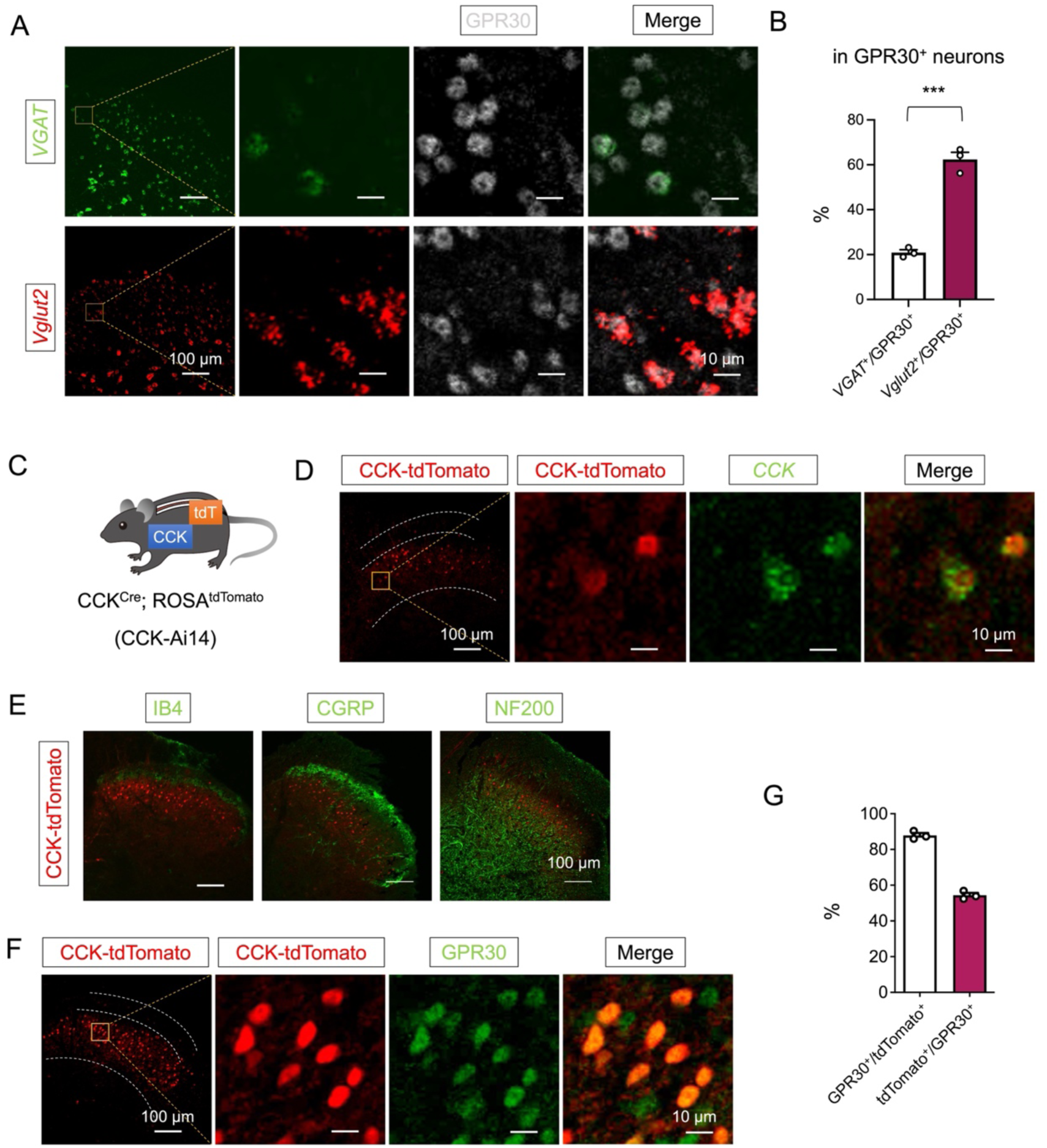
GPR30 was widely expressed in the CCK^+^ excitatory interneurons in the SDH. In situ hybridization showing *VGAT* (Green) and *Vglut2* (Red) with GPR30 (Gray) in the SDH. Scale bar: 100 μm. Boxed area of image is enlarged on the right. Scale bars: 10 μm. **B** Percentage of GPR30 positive neurons expressing *VGAT* or *Vglut2* in wild mice. (n = 3 mice, 3-4 pictures were analyzed for each mouse) **C** CCK^Cre^ mice express tdTomato in a Cre-dependent manner. **D** In situ hybridization showing CCK (Green) with tdTomato (Red) in the SDH. Scale bar: 100 μm. Boxed area of image is enlarged on the right. Scale bars: 10 μm. **E** Double staining of tdTomato^+^ neurons (Red) with IB4, CGRP or NF200 (Green) by immunochemistry. Scale bars: 100 μm. **F** Immunofluorescence in SDH to verify the co-localization of tdTomato^+^ neurons (Red) with GPR30 (Green). Scale bars: 100 μm. Boxed area of images is enlarged on the right. Scale bars: 10 μm. **G** Percentage of double labeled neurons in CCK-tdTomato and GPR30 positive neurons. (n = 3 mice, 3 pictures were analyzed for each mouse)

Given the established role of CCK^+^ neurons in neuropathic pain and their dominance among SDH excitatory interneurons^[10, 21, 27, 28]^, we hypothesized that GPR30 within this subset drives nociceptive sensitization. To validate this, we characterized CCK expression in the spinal cord via transgenic models by crossing *CCK-Cre* with *Ai14* reporter mice (Fig 2C). *Ish* and immunostaining confirmed that our transgenic mice recapitulated the majority of *CCK* expression in the deep laminae of the SDH as marked by isolectin B4 (IB4), CGRP and NF200 (Fig 2D and E). To verify the role of CCK^+^ neurons in allodynia under neuropathic pain, we performed immunostaining of c-Fos in CCK-Ai14 mice, and we confirmed that more CCK^+^ neurons were activated after CCI (Fig S4). Moreover, near-universal CCK^+^ neurons were co-localized with GPR30 (Fig 2F and G).

To functionally interrogate GPR30 and CCK^+^ neurons in neuropathic pain, we injected AAV2/9-DIO-shGper1-EGFP into lumbar SDH of *CCK-Cre* mice for conditional knock-down of *Gper1*, and 3 weeks later we subjected mice to CCI as well as subsequent behavioral tests (Fig 3A). Viral targeting accuracy was verified by immunostaining showing deep laminae-specific expression, mirroring endogenous CCK^+^ neurons distribution (Fig 3B). CCI-induced *Gper1* mRNA upregulation was significantly blunted in knockdown mice (Fig 3C, Fig S5A), confirming the transcriptional efficacy of virus.

**Figure 3.**
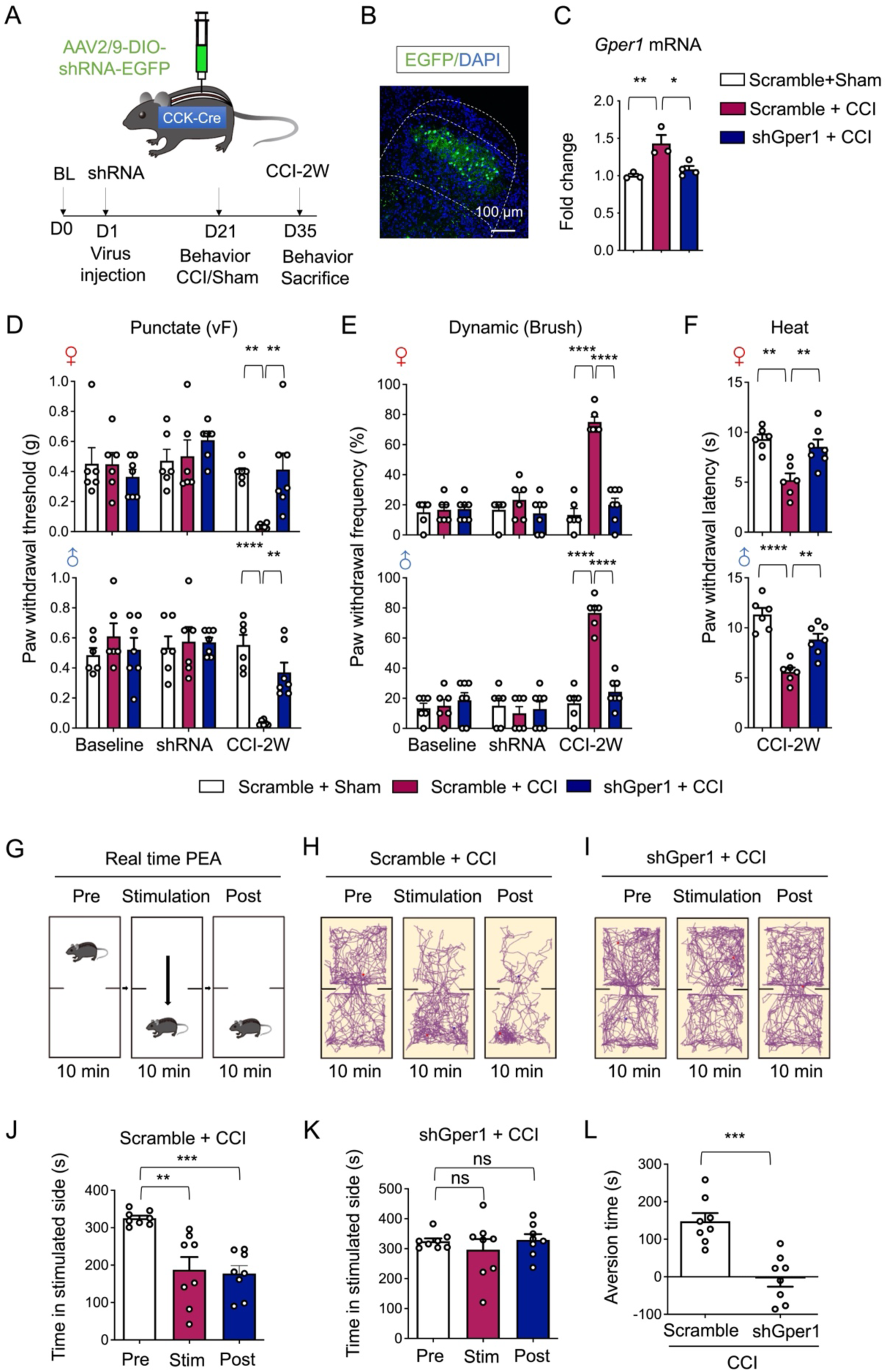
Knockdown of GPR30 in spinal CCK^+^ neurons alleviated neuropathic pain in CCI mice. **A** Schematic illustration of the strategy to knock down Gper1 in the spinal CCK^+^ neurons in a Cre-dependent manner (Up) and the diagram showing the timeline of virus injection, CCI surgery and behavioral tests (Down). **B** Immunochemical detection of the localization of virus expression (Green). Scale bars: 100 μm. **C** Quantitative PCR analysis of Gper1 mRNA in SDH from sham and CCI mice with intraspinal virus injection (n = 3-4 mice for each group). |**D-F** Behavioral tests of basic nociception, 21 days after spinal virus injection and 14 days after CCI or sham surgery along with intrathecal injection of antagonist of GPR30 or vehicle in Von Frey tests **(D)**, Brush tests **(E)** and Heat tests **(F)** in mice of both sexes (n = 6-7 mice for each group). **G** Schematic illustration of real-time place escape/avoidance (PEA) test. **H, I** Representative spatial tracking maps showing the location of a Scramble + CCI group mice **(H)** and a shGper1 + CCI group mice **(I)** before, during and after mechanical stimulation of hind paw in the chambers. **J** Quantification of time the Scramble + CCI group mice spent in preferred chamber before, during and after stimulation (n = 8 mice). **K** Quantification of time the shGper1 + CCI group mice spent in preferred chamber before, during and after stimulation (n = 8 mice). **L** Quantification of the aversion time of mice. (time spent in preferred chamber after stimulation minus the time spent before stimulation) (n = 8 mice for each group). Data information: in **(C)**, *P < 0.05; **P < 0.01 (one-way ANOVA with Turkey’s multiple comparisons test). In **(D, E)**, **P < 0.01; ****P < 0.0001 (two-way ANOVA with Turkey’s multiple comparisons test). In **(F)**, **P < 0.01; ****P < 0.0001 (one-way ANOVA with Turkey’s multiple comparisons test). In **(J, K)**, *P < 0.05; **P < 0.01; ***P < 0.001; ns = not significant. (one-way ANOVA with Turkey’s multiple comparisons test). In **(L)**, ***P < 0.001 (Unpaired Student’s t-test). All data are presented as mean ± SEM.

Behavioral tests revealed that *Gper1* knockdown in CCK^+^ neurons attenuated CCI-induced mechanical allodynia and thermal hyperalgesia across sexes, without altering basal nociceptive thresholds (Fig 3D-F). To dissect affective components of neuropathic pain, we used real-time PEA test with innocuous mechanical stimulation as shown before^[2]^ (Fig 3G). Scramble-treated CCI mice exhibited persistent aversion to stimulated side, whereas *Gper1* knockdown abolished this aversive behavior during post-stimulation period (Fig 3H-L).

### GPR30 in spinal CCK^+^ neurons is essential for the enhancement of excitatory synaptic transmission in the SDH of CCI mice

To investigate the role of GPR30 expressed on CCK^+^ neurons in neuropathic pain transmission, we conducted electrophysiological recordings using whole-cell patch clamp recording on fluorescently labeled neurons in spinal slices from CCI or sham-operated *CCK-Cre* mice treated with shRNA (Fig 4A and B). Since CCK^+^ neurons mainly receive synaptic inputs from upstream neurons, we then intended to test whether GPR30 modulated these synaptic connections. We recorded postsynaptic currents from GFP-labeled CCK^+^ neurons in laminae III-IV approximately two weeks after CCI surgery. As hypothesized, the amplitude of spontaneous excitatory postsynaptic currents (sEPSCs) in CCK^+^ neurons was significantly enhanced in CCI mice compared to sham mice, although the frequency remained unchanged (Fig 4C-E). Notably, this enhancement in amplitude was abolished in *Gper1* knockdown mice (Fig 4C, E). Notably, spontaneous inhibitory postsynaptic currents (sIPSCs) also contribute to the regulation of neuronal excitability. However, our findings indicated that knockdown of *Gper1* did not affect the IPSCs of spinal CCK^+^ neurons in CCI mice (Fig S5B-D).

**Figure 4.**
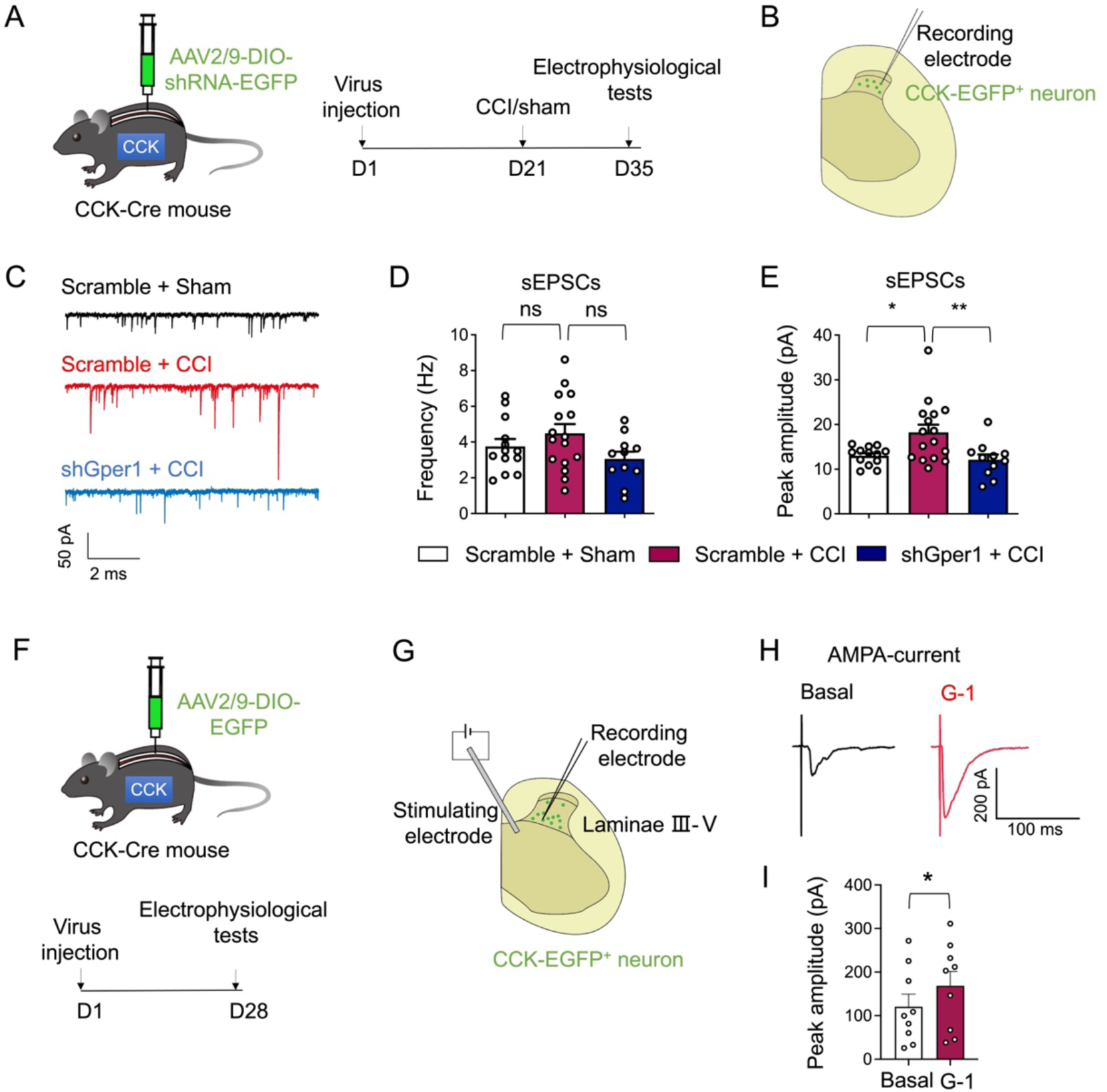
Knockdown of GPR30 in spinal CCK^+^ neurons reverse the enhancement of sEPSC in CCI mice. **A** Schematic illustration of the strategy to knock down Gper1 in the spinal CCK^+^ neurons in a Cre-dependent manner (Left) and the diagram showing the timeline of virus injection, CCI surgery and electrophysiological tests (Right). **B** Schematic illustration of spinal slice electrophysiological recordings from CCK-EGFP neurons located in the deep laminae of SDH. **C** Representative spontaneous EPSCs in CCK-EGFP neurons. **D** The frequency of spontaneous EPSCs. **E** The peak amplitude spontaneous EPSCs (n = 10-16 cells from 5 mice per group). **F** Schematic illustration of the strategy to visualize CCK^+^ neurons in a Cre-dependent manner (Up) and the diagram showing the timeline of virus injection and electrophysiological tests (Down). **G** Schematic illustration of spinal slice electrophysiological recordings from CCK-EGFP^+^ neurons located in the deep laminae of SDH during electric stimulation of the deep laminae of SDH to simulate APMA currents. **H** Representative AMPA-current in CCK-EGFP^+^ neurons before and after administration of G-1 (0.1 μM). **I** The peak amplitude AMPA currents before and after administration of G-1 (n = 8 cells from 4 mice). Data information: in **(D, E)**, *P < 0.05; **P < 0.01; ns = not significant (one-way ANOVA with Turkey’s multiple comparisons test). In **(I)**, *P < 0.05 (Paired Student’s t-test). All data are presented as mean ± SEM.

Given that EPSCs are primarily mediated through glutamatergic receptors such as AMPA receptor, coupled with emerging evidence showing that GPR30 could facilitate excitatory transmission via increasing the clustering of glutamatergic receptor subunits^[25]^, we proceeded to examine whether GPR30 activation modulated AMPA-mediated currents. Electrophysiological experiments were carried out in spinal slices from *CCK-Cre* mice treated with intraspinal injections of AAV2/9-DIO-EGFP (Fig 4F). To record AMPA-dependent currents, electronical stimulation from the electrode placed in the deep laminae of SDH was applied (Fig 4G). And CCK^+^ neurons were held at -70 mV in the presence of APV (100 μM) to block NMDA receptors and bicuculline (20 μM) to block GABA_A_ receptors and strychnine (0.5 μM) to block glycine receptors. All recorded cells responded to the electrode stimulation, exhibiting evoked EPSCs (Fig 4H and I). Furthermore, the application of 0.1 μM G-1 augmented the amplitude of AMPA currents (Fig 4H and I). These findings collectively suggest that GPR30 regulates sEPSCs in CCK^+^ neurons through an AMPA-dependent mechanism.

### GPR30 is expressed in the spinal CCK^+^ neurons receiving direct projection from S1 cortex

To anatomically verify that the lumbar spinal cord receives direct descending projections from the S1 cortex, we employed a combination of retrograde and anterograde tracing methods to investigate the connections between cortical neurons and their axonal innervation in the SDH. Our results revealed that retrograde labeling with cholera toxin subunit B-555 (CTB-555), injected into the deep laminae of the lumbar SDH, predominantly localized neurons in the contralateral S1 cortex (Fig 5A-C). Of note, the neurons projecting from S1 cortex are primarily excitatory pyramidal neurons located in layer V^[29]^. Similarly, fiber tracing using AAV2/9-hSyn-EGFP indicated that axons from S1 terminate extensively in laminae III-V of the contralateral SDH (Fig 5D-F).

**Figure 5.**
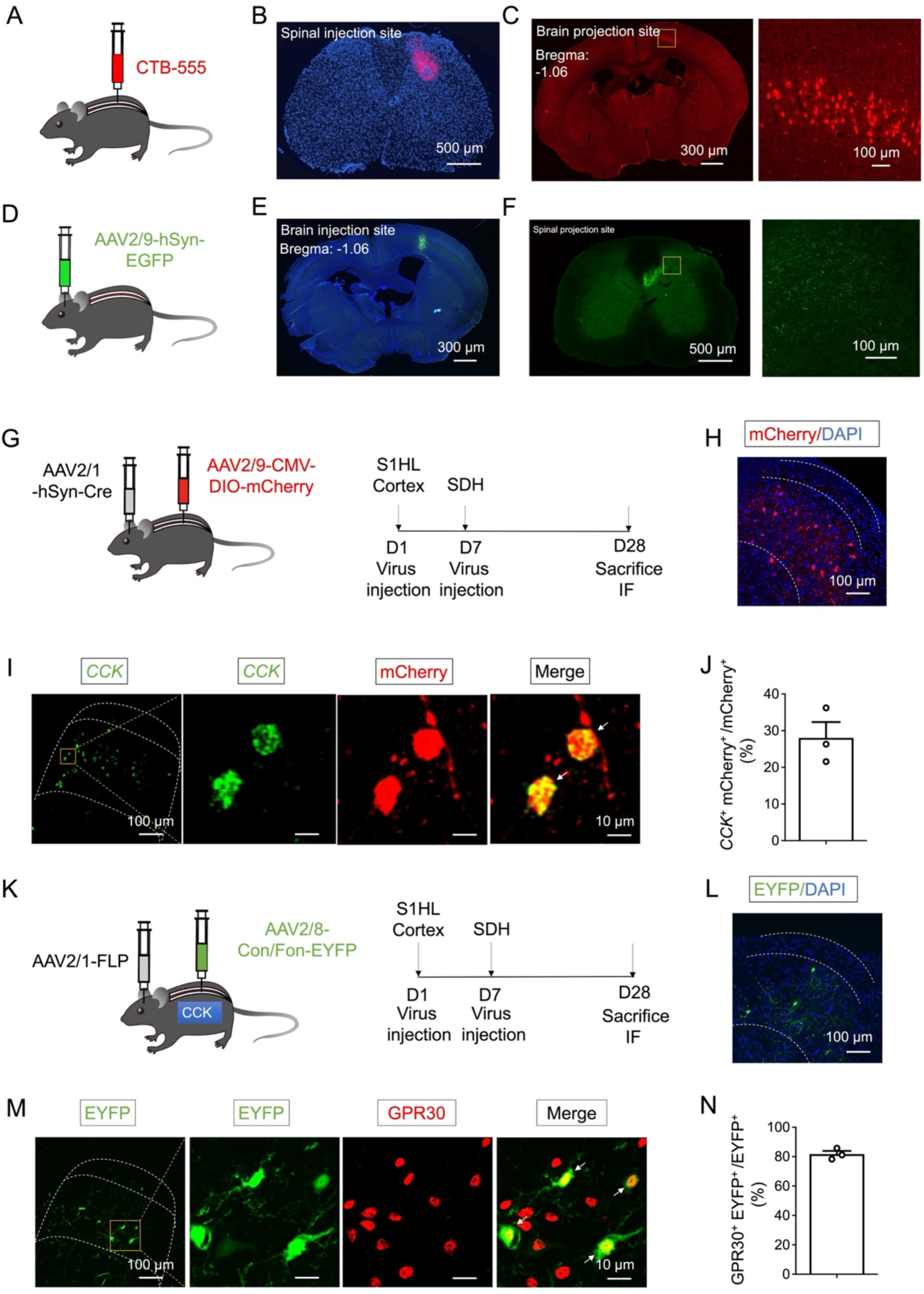
GPR30 was expressed in the spinal CCK^+^ neurons receiving direct projection from S1 cortex. **A** Schematic illustration of the strategy to retrograde tracing the projections innervating spinal dorsal horn via CTB-555. **B** Fluorescence image showing the site of CTB-555 injection in the spinal dorsal horn. Scale bars: 500 μm. **C** Coronal brain section showing the location of tdTomato+ neurons in the S1HL cortex. Scale bars: 300 μm. Boxed area of image is enlarged on the right. Scale bars: 100 μm. **D** Schematic illustration of the strategy to anterograde tracing the projections from S1HL cortex. **E** Coronal brain section showing the site of AAV injection in the S1HL cortex. Scale bars: 300 μm. **F** Fluorescence image showing the S1HL-SDH tract terminates in the deep laminae of spinal dorsal horn. Scale bars: 500 μm. Boxed area of image is enlarged on the right. Scale bars: 100 μm. **G** Schematic illustration of the strategy for identifying the spinal post-synaptic neurons of S1-SDH projections (Left) and diagram showing the timeline of AAV2/1 injection in the S1HL cortex, AAV2/9 injection in the spinal dorsal horn and immunofluorescence (Right). **H** Representative image showing the mCherry^+^ post-synaptic neurons of the S1HL-SDH projections in the deep laminae of SDH. Scale bars: 100 μm. **I** In situ hybridization showing CCK (Green) with RFP (Red) by immunofluorescence in the spinal dorsal horn. Scale bars: 100 μm. Boxed area of image is enlarged on the right. Scale bars: 10 μm. White arrows indicate double-positive cells. **J** About 28.13% of RFP^+^ post-synaptic neurons of S1HL-SDH projections expressing CCK (n = 3 mice, 3 pictures were analyzed for each mouse). **K** Schematic illustration of the strategy for identifying the spinal CCK^+^ post-synaptic neurons of S1HL-SDH projections in CCK^Cre^ mice (Left) and diagram showing the timeline of AAV2/1 injection in the S1HL cortex, AAV2/9 injection in the spinal dorsal horn and immunofluorescence (Right). **L** Representative image showing the EGFP^+^ CCK^+^ post-synaptic neurons of the S1-SDH projections in the spinal dorsal horn with DAPI. Scale bars: 100 μm. **M** Double staining of EGFP^+^ neurons with GPR30 by immunohistochemistry. Scale bars: 100 μm. Boxed area of image is enlarged on the right. Scale bars: 10 μm. **N** About 81.7% of CCK+ post-synaptic neurons of S1HL-SDH projections expressing GPR30 (n = 3 mice, 7-18 pictures were analyzed for each mouse). Data information: in **(J, N)**, data are presented as mean ± SEM.

Building on previous findings suggesting a functional interaction between S1-SDH projections and spinal CCK^+^ neurons^[13]^, our current study aimed to further elucidate the structural relationship among GPR30, S1-SDH projections and CCK^+^ neurons. To achieve this, we performed anterograde trans-monosynaptic tracing by injecting AAV2/1-hSyn-Cre into the right S1 cortex, followed by AAV2/9-DIO-mCherry into the contralateral lumbar SDH. This approach allowed us to visualize the mCherry^+^ S1-SDH postsynaptic neurons in the deep laminae of the lumbar SDH (Fig 5G and H). Consistent with prior observations, we found that 28.1% of mCherry+ S1-SDH downstream neurons coexpressed CCK (Fig 5I and J). Thus, our findings confirm that spinal CCK^+^ neurons are innervated by the S1 cortex.

To further examine the interplay between GPR30, S1-SDH projections, and CCK^+^ neurons, we utilized a monosynaptic anterograde tracing strategy. Specifically, we injected AAV2/1-EF1α-FLP into the right S1 cortex and AAV2/9-hSyn-Con/Fon-GFP into the contralateral lumbar SDH of CCK-Cre mice. This strategy enabled us to visualize GFP-positive S1-SDH postsynaptic CCK^+^ neurons (Fig. 5K and L). Our co-staining results revealed that the vast majority of CCK^+^ S1-SDH postsynaptic neurons expressed GPR30 (Fig 5M and N). These data collectively indicate that the majority of CCK^+^ neurons receiving S1 projections express GPR30.

### GPR30 in S1-SDH post-synaptic neurons is critical for CCI-induced neuropathic pain

Given that GPR30 has been verified to be expressed on CCK^+^ neurons receiving S1-SDH direct projections (Fig 5M), we employed a combination of chemogenetic and pharmacological approaches to determine whether neurons innervated by S1-SDH direct projections mediate nociception via GPR30^[30]^. Specifically, we performed injections of anterograde AAV2/1-hSyn-Cre into the right S1 WT mice. One week later, we administered AAV2/9-hSyn-DIO-hM3Dq (Gq)-mCherry into the lumbar SDH (Fig 6A). Pharmacological activation of these post-synaptic neurons with CNO in Gq-treated mice significantly induced spontaneous pain-like behaviors, such as paw scratching, biting, and licking, in the hind paws and tails (Fig 6B). Furthermore, chemogenetic activation of S1-SDH post-synaptic neurons dramatically induced mechanical allodynia and thermal hyperalgesia in both sexes compared to negative control groups (Fig 6C-E). Notably, the reduction in nociceptive thresholds could be effectively reversed by intrathecal administration of G-15 (Fig 6C-E). The corresponding immunohistochemistry results using c-Fos confirmed the chemogenetic activation of S1-SDH post-synaptic neurons, which could be suppressed by intrathecal application of G-15 (Fig 6F-H).

**Figure 6.**
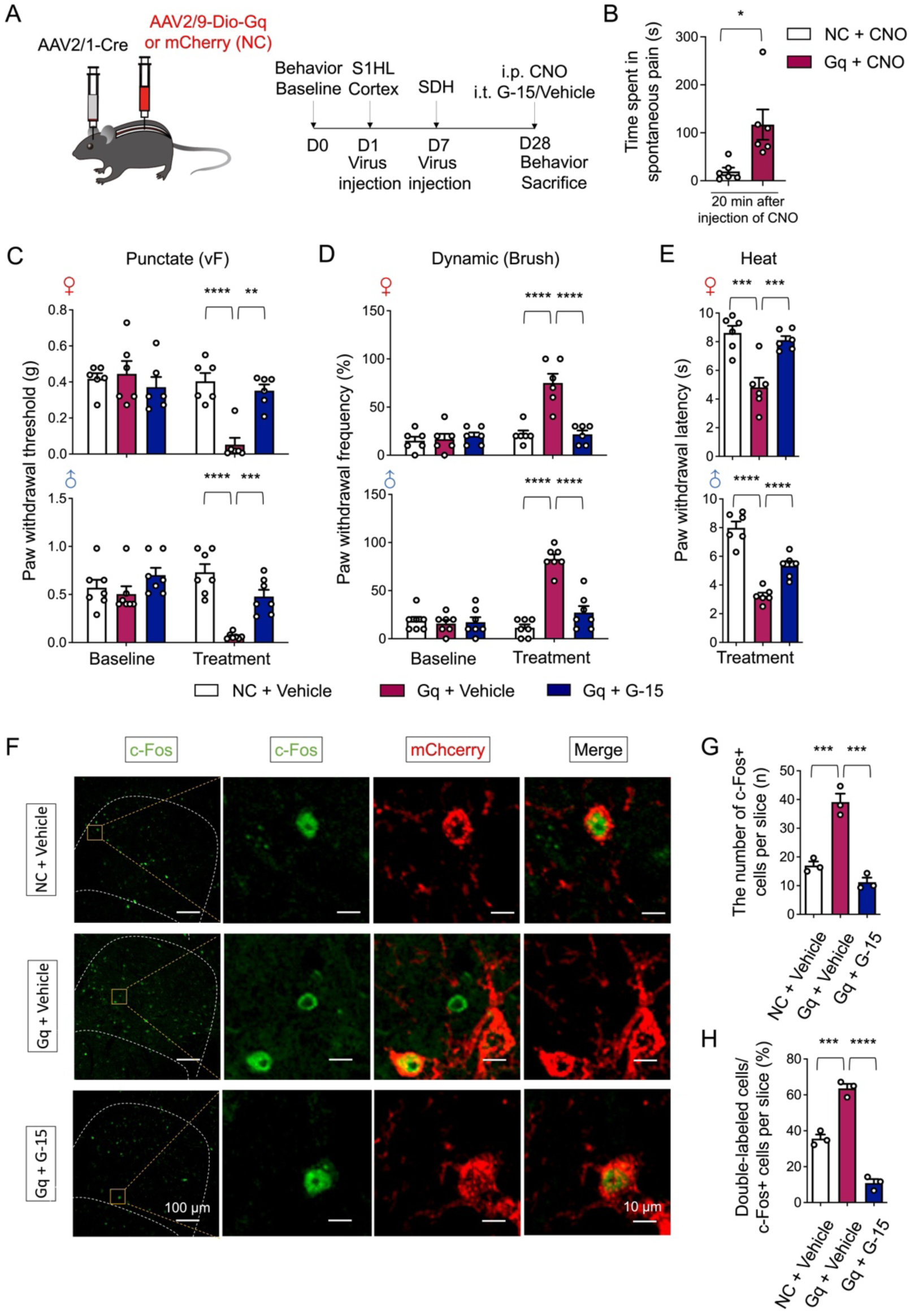
Chemogenetic activation of S1-SDH post-synaptic neurons mimicked neuropathic pain symptoms, which were reversed by spinal inhibition of GPR30. **A** Schematic illustration of the strategy for identifying the post-synaptic neurons of S1HL-SDH projections in SDH (Left) and diagram showing the timeline of AAV2/1 injection in the S1HL cortex, AAV2/9 injection in the spinal dorsal horn and behavioral tests (Right). **B** Spontaneous pain induced by intraperitoneal injection of CNO within 20 min (n = 6 mice for each group). **C-E** Behavioral tests of basic nociception, 28 days after brain virus injection along with intraperitoneal injection of CNO and intrathecal injection of antagonist of GPR30 or vehicle in Von Frey tests **(C)**, Brush tests **(D)** and Heat tests **(E)** in mice of both sexes (n = 6 mice for each group). **F** Immunochemical detection of c-Fos (Green) and m-Cherry+ post-synaptic neurons (Red). Scale bars: 100 μm. Boxed area of images is enlarged on the right. Scale bars: 10 μm. **G** Total number of c-Fos positive neurons in the SDH per section (n = 3 mice for each group, 3 pictures were analyzed for each mouse). **H** Percentage of c-Fos positive neurons expressed in m-Cherry (n = 3 mice for each group, 3 pictures were analyzed for each mouse). Data information: in **(B)**, *P < 0.05 (Unpaired Student’s t-test). In **(C, D)**, **P < 0.01; ***P < 0.001; ****P < 0.0001 (two-way ANOVA with Turkey’s multiple comparisons test). In **(E)**, ***P < 0.001; ****P < 0.0001 (one-way ANOVA with Turkey’s multiple comparisons test). In **(G, H)**, ***P < 0.001; ****P < 0.0001. (one-way ANOVA with Turkey’s multiple comparisons test). All data are presented as mean ± SEM.

To further explore the role of S1-SDH post-synaptic neurons in the modulation of neuropathic pain, we employed chemogenetic inhibitory methods to suppress these neurons in CCI mice (Fig S6A). The suppression of S1-SDH post-synaptic neurons could dramatically relieve the mechanical allodynia and thermal hyperalgesia induced by CCI (Fig S6B-D). To establish the essential role of GPR30 in this process, we specifically knocked down the expression of *Gper1* on S1-SDH post-synaptic neurons and subjected mice to CCI after adequate viral expression (Fig 7A). Interestingly, the knockdown of *Gper1* in S1-SDH post-synaptic neurons was sufficient to relieve mechanical allodynia and thermal hyperalgesia in both sexes (Fig 7B-D). Immunochemistry showed the viral location in deep laminae of SDH (Fig 7E) and qPCR confirmed the suppression of *Gper1* mRNA expression which was increased by CCI (Fig 7F). Collectively, these findings underscore the role of GPR30 in the descending facilitation potentially mediated by S1-SDH projections in the neuropathic pain.

**Figure 7.**
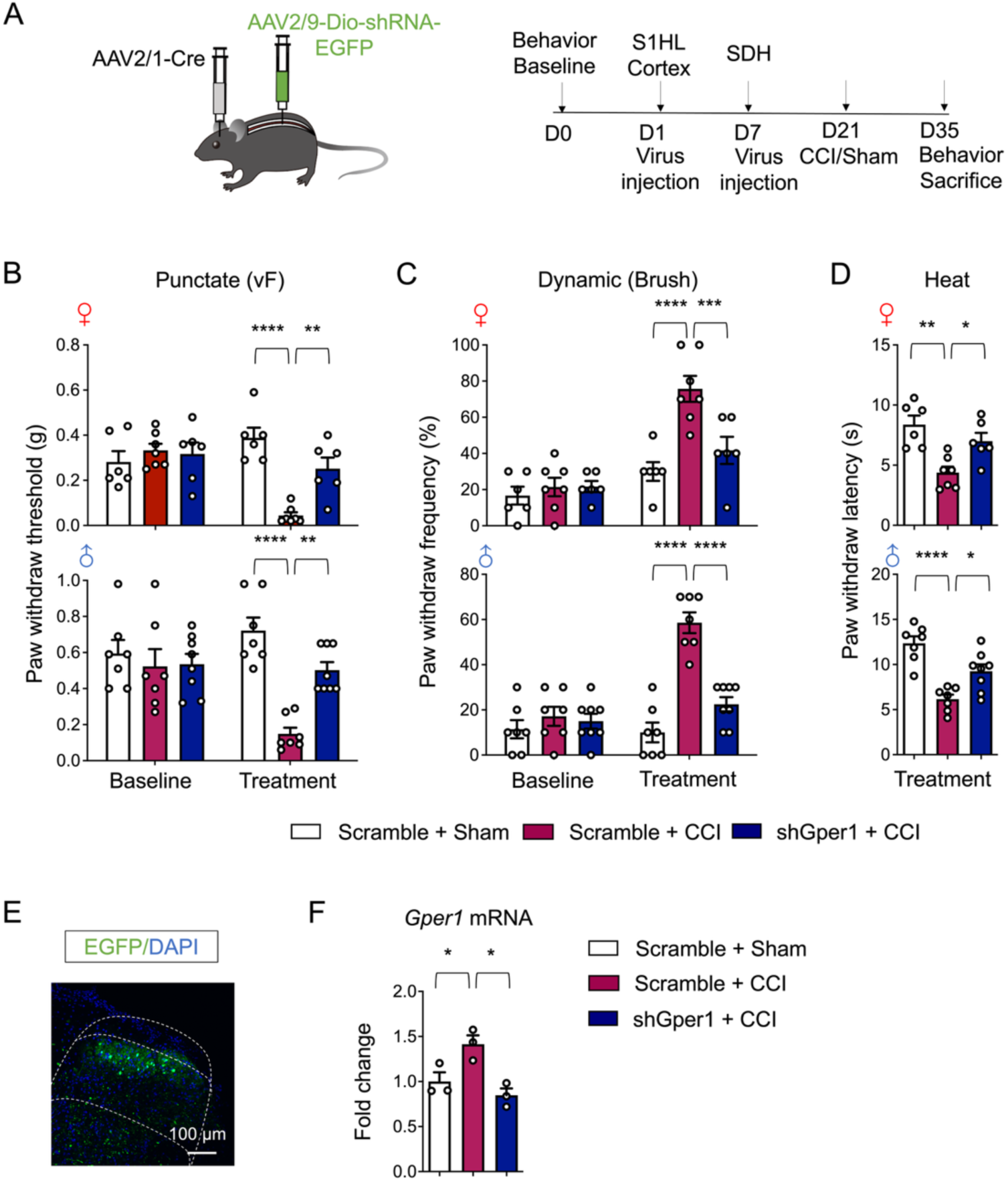
CCI-induced neuropathic pain was attenuated by knock-down of GPR30 in S1-SDH post-synaptic neurons. **A** Schematic illustration of the strategy to knock down Gper1 in the post-synaptic neurons of S1HL-SDH projections in a Cre-dependent manner (Up) and the diagram showing the timeline of virus injection, CCI surgery and behavioral tests (Down). **B-D** Behavioral tests of basic nociception and 14 days after CCI surgery in Von Frey tests **(B)**, Brush tests **(C)** and Heat tests **(D)** in mice of both sexes (n = 6 mice for each group). **E** Immunochemical detection of the localization of virus expression (Green). Scale bars: 100 μm. **F** Quantitative PCR analysis of Gper1 mRNA in SDH (n = 3 mice for each group). Data information: in **(B, C)**, **P < 0.01; ***P < 0.001; ****P < 0.0001 (two-way ANOVA with Turkey’s multiple comparisons test). In **(D)**, *P < 0.05; **P < 0.01; ****P < 0.0001 (one-way ANOVA with Turkey’s multiple comparisons test). In **(F)**, *P < 0.05 (one-way ANOVA with Turkey’s multiple comparisons test). All data are presented as mean ± SEM.

## Discussion

CCK^+^ neurons, located in the deep laminae of the spinal cord, have long been recognized for their pivotal role in the development and maintenance of neuropathic pain, as well as descending facilitation by sensory cortex-spinal cord projections. Despite extensive research, the molecular mechanisms underlying nociception remain poorly understood. GPR30, as a membrane estrogen receptor, exerts modulatory effects on various physiological and pathological processes, including the development of neuropathic pain, within seconds to minutes ^[22]^. In this study, we observed significant upregulation of GPR30 expression in the SDH after CCI, underscoring its critical role in neuropathic pain development. To investigate this further, we employed a comprehensive approach, including transgenic mice models, behavioral assays, pharmacological interventions, chemogenetic methods, and electrophysiological studies. These experiments revealed that GPR30 plays an indispensable role in neuropathic pain, particularly through its regulation of AMPA-dependent EPSCs in spinal CCK^+^ neurons, which are essential for pain signaling. Furthermore, we demonstrated that GPR30 expressed on post-synaptic neurons of corticospinal direct projections took a role in modulation of neuropathic pain. These findings collectively suggest that GPR30 in spinal CCK^+^ neurons may represent a promising therapeutic target for neuropathic pain.

Estrogen has long been recognized as a key player in modulating nociception, with its drastic fluctuations notably impacting nociceptive thresholds^[31]^. Our prior research demonstrated that moderate estrogen supplementation effectively mitigated hyperalgesia in ovariectomized (OVX) mice, however, excessive estrogen supplementation paradoxically exacerbated hyperalgesia^[31]^. This discrepancy might stem from the fact that varying estrogen concentrations can differentially activate estrogen receptors, including nuclear receptors (ERα and ERβ) and membrane receptors (GPR30)^[22]^. Considering our findings that inhibiting spinal GPR30 does not alter the basal nociception in naïve mice, it appears that GPR30 may not be significantly activated by estrogen under normal conditions. Additionally, estrogen has long been implicated in modulating neuropathic pain. Intra-dorsal root ganglion (DRG) administration of estrogen in CCI rats has been shown to enhance mechanical and thermal pain in an ERα-dependent manner^[32]^. Conversely, estrogen supplementation in the anterior cingulate cortex (ACC) of CCI mice significantly alleviated neuropathic hyperalgesia in a GPR30-dependent manner^[33]^. These findings further underscore the distinct roles of different estrogen receptors in pain modulation.

GPR30 is widely expressed in the nervous system and exerts vital effects in nociceptive modulation^[16, 23, 24, 33–37]^. For example, the activation of GPR30 in DRG could aggravate the hyperalgesia in OVX mice, while inhibition of GPR30 relieved hyperalgesia^[38]^. Besides, the GPR30 expressed on GABAergic cells in rostral ventromedial medulla (RVM) mediates the descending facilitation of nociception^[24]^. However, though GPR30 is also widely expressed in SDH^[26]^, still little is known about the functions as well as underlying mechanisms of spinal GPR30 in nociceptive modulation. Consistent with previous studies^[39]^, here we found that intrathecal injection of G-1 could dramatically induce mechanical allodynia and thermal hyperalgesia in mice. To further explore whether spinal GPR30 is involved in pathological nociception, we subjected mice to CCI surgery to mimic neuropathic pain^[2, 40]^. In accordance with our expectations, the inhibition of spinal GPR30 significantly reversed the mechanical allodynia and thermal hyperalgesia induced by CCI. Moreover, GPR30 has been reported to be involved in nociceptive sexual dimorphism. For example, the regulatory role of GPR30 in DRG in maintenance of hyperalgesia induced by repeated exposure of opioid only exists in female rats^[41, 42]^. However, according to our results, spinal GPR30 modulated nociception in a sex-independent manner. We also found that GPR30 is indiscriminately expressed on the neurons of both sexes of mice, which might account for the sex-independent function of spinal GPR30 in nociceptive modulation. Consistent with our results, several studies also have confirmed that the spinal *Gper1* expression showed no significant difference between male and female mice^[43]^. Furthermore, the fluctuation of estrogen failed to change the basic expression of spinal GPR30^[31]^. These results further indicate that spinal GPR30 modulated nociception in a sex-independent manner.

The SDH is a major locus for the integration of peripheral sensory input and supraspinal modulation. Most peripheral nociceptive afferents project to the superficial laminae of the SDH which respond to the noxious stimulations, while low-threshold mechanoreceptors form synaptic contacts in the deep laminae which respond to innocuous stimulations^[8]^. However, under the condition of mechanical allodynia, innocuous stimulation might also activate more superficial nociceptive circuits and lead to painful perception, which might come from the circuit-based transformation in the SDH^[6, 44]^. It should be noted that the spinal dorsal horn is composed of a large number of excitatory (75%) and inhibitory (25%) interneurons, as well as a small part of projection neurons which relay integrated information to various supraspinal regions^[9]^. Excitatory interneurons have been confirmed to take a vital role in conveying mechanical allodynia according to the nature of injury^[10, 45]^. As a distinct type of excitatory interneurons mainly located in the deeper laminae of SDH, CCK^+^ neurons are important for neuropathic injuries^[10–12]^. The inhibition of spinal CCK^+^ neurons could alleviate neuropathic mechanical allodynia to a great extent^[10]^.In addition, spinal CCK^+^ neurons also account for the thermal hyperalgesia^[12]^. However, little is known how CCK^+^ neurons mediate the nociception. Combined with our results that GPR30 is widely expressed in spinal excitatory interneurons and involved in neuropathic pain modulation, we speculate that CCK^+^ neurons might convey neuropathic hyperalgesia via GPR30. As expected, most CCK^+^ neurons express GPR30 and knock-down of the *Gper1* in CCK^+^ neurons dramatically relieve the pain induced by CCI, thus indicating the vital role of GPR30 in CCK^+^ neurons mediating neuropathic pain. However, it should be noted that half of GPR30^+^ neurons are not co-localized with CCK^+^ neurons, and further studies are needed to explore the function of these GPR30^+^CCK^-^ neurons in neuropathic pain.

Abnormal activation of neurons in SDH is one of the causes of hyperalgesia and the change of post synaptic currents is the vital factor influencing neuronal excitability^[46, 47]^. Given this critical link between EPSCs and excitability, we measured excitatory postsynaptic currents (EPSCs) and found an elevation of EPSCs amplitudes in spinal CCk^+^ neurons after CCI. Furthermore, the knock-down of *Gper1* in CCK^+^ neurons could inhibit the increase of EPSCs amplitude induced by CCI. Together, these data illustrate that GPR30 promotes the enhancement of synaptic transmission. It should be noted that EPSCs are specifically produced by glutamatergic receptors expressed on post-synaptic membrane, including AMPA and NMDA receptors^[25, 46, 48]^. In our study, we confirmed that the selective activation of GPR30 by G-1 remarkedly enhanced the AMPA-current in spinal CCK^+^ neurons, which might account for the increased excitability of CCK^+^ neurons in neuropathic pain. It should be noted that the IPSCs could also influence the excitability of neurons, however, the knockdown of Gper1 failed to change the IPSCs amplitude in CCI mice, suggesting that GPR30 did not take part in the inhibitory synaptic regulation. However, it should be noted that our data did not rule out the potential pre-synaptic contributions.

Increasing evidence has mapped neural circuits from peripheral to central nervous system to illustrate the neural mechanisms of nociception^[5, 7–9, 45, 49]^. In brief, pain is derived from the activation of peripheral nociceptors whose cell bodies lie in DRG, and then nociceptive signals are transduced to the SDH for preliminary regulation and finally projected to cerebral cortex via a series of brain region mediating nociception. Additionally, pain is also modulated by the descending modulatory pathways constituted of projections from ventrolateral periaqueductal gray (PAG) to the RVM and then to the spinal cord which takes several steps^[50–55]^. However, A recent study has come up with the existence of long direct projections form S1 cortex to deep laminae of SDH and the vital role of S1-SDH projections in neuropathic pain^[13]^. Inhibition of S1-SDH projections attenuates neuropathic pain, while activation decreases pain thresholds in naive mice. We confirmed the existence of these direct projections and their postsynaptic targets’ critical role in neuropathic pain. Notably, these long projections specifically synapse within SDH deep laminae onto CCK⁺ neurons^[13]^.We also structurally verify that CCK^+^ neurons receive projections from S1 cortex. Furthermore, we also found that the majority of CCK^+^ neurons receiving S1-SDH projections express GPR30, thus indicating an important role of GPR30 in descending modulation of S1-SDH. As expected, the knockdown of *Gper1* in S1-SDH post-synaptic neurons dramatically alleviated the hyperalgesia induced by CCI. All these results further suggest an important role of GPR30 in descending facilitation of neuropathic pain. However, since viral strategy only labeled a small fraction of post-synaptic CCK^+^ neurons from S1 cortex and was insufficient to functionally manipulate these neurons, more efficient methods should be employed to verify the role of GPR30 expressed on S1-SDH post-synaptic CCK^+^ neurons under neuropathic pain conditions. In addition, manipulation of the S1-SDH projections should be employed in the future to further verify the direct functional connection between corticospinal projections and GPR30.

## Conclusion

GPR30 plays a critical role in the development of neuropathic pain, particularly within CCK+ neurons. Our research highlights GPR30’s essential function in enhancing AMPA-dependent EPSCs, which are crucial for the activation of CCK^+^ neurons and the subsequent development of abnormal nociception under neuropathic conditions. Furthermore, GPR30 in post-synaptic neurons of the descending projections via corticospinal projections contributes to the propagation of neuropathic pain signals. These findings suggest that targeting GPR30 in spinal CCK^+^ neurons could be a promising therapeutic strategy for neuropathic pain in clinic.

## Materials and methods

### Animals

Mice of both sexes ranging in age from 8 weeks to 12 weeks were used for this study, including C57BL/6JRJ wild-type (purchased from SLAC Laboratory Animal CO. LTD, Shanghai, China), *CCK-Cre* mice and *Ai14* mice (originally purchased from Jackson Laboratory). In accordance with the Jackson Laboratory’s protocol, transgenic mice were genotyped. All animals were kept in a humidity-controlled room with free access to food and water, the facility was maintained at 22 ℃ and ran on 12 hours of light/dark cycles. A random assignment of animals to different experiment groups was conducted. The animals were treated in accordance with protocols approved by the Animal Ethic and Welfare Committee of Zhejiang University School of Medicine, and all experimental procedures were carried out in accordance with the National Institute of Health Guide for Care and Use of Laboratory Animals (NIH Publications NO.86-23).

### Drug administration

For pharmacological manipulation of the activity of spinal GPR30, G-1 or G-15 (diluent of 0.2 mg/mL, administration of 100 μg/kg, 10 μL per mice; APExBIO, USA) was dissolved in 1% DMSO with Saline and administered intrathecally as previously described^[39]^. To be specific, mice were lightly anesthetized with 1.5% inhaled isoflurane, and held with a pen under the pelvis while a 25-gauge needle attached to a 10-μL syringe (Hamilton, Nevada, USA) was inserted in the subarachnoid space between vertebrae L5 and L6 until a tail flick was observed. The syringe was held for 30 seconds after the injection of 10 μL solution per mice. For chemogenetic manipulation of S1-SDH post-synaptic neurons, Clozapine N-oxide (CNO; 2.5 mg/kg, 150 μL per mice; MCE, China) was dissolved in saline with gentle vortex for mixing and then administered intraperitoneally ^[30]^. The behavioral assessments were carried out 30 minutes following the injection.

### Virus and CTB microinjection

For intracranial injection, mice were anesthetized with 1% pentobarbital sodium solution (70 mg/kg per mice) and then secured in a stereotaxic frame (RWD Life Science, Shenzhen, China). A middle scalp incision exposed the skull, and then a hole was drilled on the skull above the right S1 cortex to allow passage of a glass microelectrode filled with the virus. Viral injections were performed with the following coordinates of S1: 0.95∼1.15 mm from bregma, 1.4∼1.6 mm from midline, and 0.9∼1.1 mm ventral to skull. A volume of 300 nL virus was injected at 50 nL/min with calibrated glass microelectrodes by a microsyringe pump (#78-8710 KD Scientific, USA). After infusion, the micropipette was slowly removed after five minutes. For spinal cord injection, mice were anesthetized with 1% pentobarbital sodium solution. The spinal cord could be visible between T12 and T13 vertebral spines following a middle incision along the lumbar vertebrae. With a stereotaxic frame, a glass microelectrode was inserted between L3-L4 spinal cord to a depth of -400 um below the dura, avoiding the posterior spinal arteries. With a stereotaxic injector, 500 nL of viral solution or CTB-555 (1% in PBS; BrainVTA, Wuhan, China) was slowly infused over a period of 5 minutes. The micropipette was left in place for 5 minutes after infusion before being slowly removed.

For knock-down the *Gper1* in CCK^+^ neurons, AAV2/9-CMV-DIO-(EGFP-U6)-shRNA (GPR30)-WPRE-pA (5×10^12^ v.g./mL) or AAV2/9-CMV-DIO-(EGFP-U6)-shRNA (Scramble)-WPRE-pA (5×10^12^ v.g./mL) was injected into the lumbar SDH of *CCK-Cre* mice. For visualization of the CCK+ neurons, AAV2/9-CMV-DIO-EGFP-WPRE-pA (5.2×10^12^ v.g./mL) was injected into the lumbar SDH of *CCK-Cre* mice. For anterograde tracing of S1 cortex projections, AAV2/9-hSyn-EGFP-WPRE (1×10^13^ v.g./mL) was in injected into the S1 cortex of wild type mice. For visualization of the S1-SDH post-synaptic neurons, AAV2/1-hSyn-CRE-WPRE-pA (1×10^13^ v.g./mL) was injected into the S1 cortex and AAV2/9-EF1α-DIO-mCherry-WPRE-pA (1×10^13^ v.g./mL) was injected into the lumbar SDH in wild type mice. For visualization of CCK^+^ post-synaptic neurons of S1-SDH projections, AAV2/1-EF1α-FLP-WPRE-pA (1×10^13^ v.g./mL) was injected into the S1 cortex and AAV2/8-hSyn-Con/Fon-EYFP-WPRE-pA (2×10^12^ v.g./mL) was injected into the lumbar SDH of *CCK-Cre* mice. For chemogenetic manipulation of S1-SDH post-synaptic neurons, AAV2/1-hSyn-CRE-WPRE-pA (1×10^13^ v.g./mL) was injected into the S1 cortex, while AAV2/9-hSyn-DIO-hM3Dq (Gq)-mCherry (3.3×10^13^ v.g./mL; dilution: 1:5) or AAV2/9-hSyn-DIO-hM4Di (Gi)-mCherry (3.3×10^13^ v.g./mL; dilution: 1:5) or AAV2/9-hSyn-DIO-mCherry (3×10^13^ v.g./mL; dilution: 1:5) was injected into the lumbar SDH in wild type mice. All viruses mentioned above were purchased from BrainVTA (Wuhan, China). For visualization of the localization of excitatory interneurons in the SDH, AAV2/9-hSyn-mCaMkⅡa-mCherry-WPRE-pA (1×10^13^ v.g./mL; Taitool Bioscience, Shanghai, China) was injected into the lumbar SDH of wild type mice.

For structural tests, at least 3 mice were examined and each mouse was examined at least 3 slices. For behavioral tests, at least 6 mice per group were examined. The mice with improper position or expression of virus were excluded.

### Chronic constriction injury (CCI)

The CCI-induced neuropathic pain model was employed as previously documented ^[2]^, Mice were lightly anesthetized via inhaled 1.5% Isoflurane. An incision was made on the skin of each mouse, exposing the sciatic nerve. Four ligations with 6-0 chromic silk were loosely tied around the sciatic nerve. Nerve constriction should be minimal until a brief twitch can be observed. In sham mice, the sciatic nerve was exposed without ligation. The animal was allowed to recover from surgery for 2 weeks before behavioral testing.

### Behavioral test

#### Punctate mechanical stimuli (von Frey filaments)

Mice were habituated to opaque cage (7.5×15×15 cm) for 1 hour the day before and 30min immediately prior to testing. Testing was performed using a series of von Frey filaments using the Dixon’s Up-down method^[56]^, beginning with the 0.16 g filament. The 50% paw withdrawal threshold was determined for each mouse on one hind paws. Each filament was gently applied to the plantar surface of the hind paw for 5 seconds or until a response such as a sharp withdraw, shaking or licking of the limb was observed. Between individual measurement, filaments were applied at least 3 minutes after the mice had returned to their initial resting state.

#### Dynamic mechanical stimuli (Brush)

Each mouse was habituated in an opaque cage (7.5×15×15 cm) for 1 hour the day before and 30 minutes immediately prior to testing. The plantar hind paw was stimulated by light stroking from heel to toe with a paintbrush. A positive response was recorded if the animal lifting, shaking or licking the limb. The application was repeated 10 times with a 3 minutes interval between each stimulation.

#### Plantar heat test (Hargreaves Method)

Mice were placed in an acrylic chamber on a glass table and allowed to acclimate to the test chamber for 1hour the day before and 30 minutes immediately prior to testing. The thermal paw withdrawal latency was assessed using the plantar test (Ugo Basile Biological Research Apparatus, Gemonio, Italy). While the mouse was in a motionless state, a radiant heat source, which was maintained at 40 W, was applied to the plantar surface of the mouse’s paw through the glass plate. The paw withdrawal latency was defined as the time to withdrawal of the hind paw from the heat source, and 15 seconds was used as the cut-off to avoid injury.

#### Real-time place escape/avoidance test (RT-PEA)

Each mouse was habituated in the test room for 1 hour the day before and 30 minutes immediately prior to testing. The Real-time place escape/avoidance chamber (50×28×32 cm; made with plastic plates that had distinct color with another) was placed on the mesh floor. The tested mouse was placed in a two-chamber box and allowed to explore both chambers without any stimulation (pre-stimulation, 10 minutes); mechanical simulation by 0.16 g von Frey filament was intermittently delivered whenever the mouse entered or stayed in the preferred chamber, as shown in the pre-stimulation stage (stimulation, 10 minutes); the mouse then freely explored the box without any stimulation (post-stimulation, 10 minutes). The mouse’s movements and time stay in preferred chamber were recorded via an ANY-Maze system.

### Immunohistochemistry

Mice were deeply anesthetized with 1% pentobarbital sodium solution and then perfused with phosphate-buffered saline (PBS) followed by pre-cooled 4% paraformaldehyde fix solution (PFA). For c-Fos staining, Von Frey filament with the same force (0.16 g) representing light mechanical stimulation was applied to the right hind paw of each group every 30 s for 20 min. Then animals were then perfused with PBS and 4% PFA ninety minutes after Von Frey filament stimulation. Tissues were harvested and post-fixed in PFA at 4℃ over night before being dehydrated in 30% sucrose for 2days. Tissues were embedded in Optimal Cutting temperature (OCT) and then cut into 10-30 μm sections placed directly onto slides. Tissue slices were blocked at room temperature for an hour with block solution containing 10% normal donkey serum (NDS), 1% bovine serum albumin BSA and 0.3% triton X-100 in PBS (PBS-T), and then incubated with primary antibodies diluted in 1% NDS, 1% BSA in PBS-T at 4℃ overnight. Sections were washed in PBS and incubated with secondary antibodies at room temperature for 1-2 hours. Slices were washed and covered with Fluoromount-G containing DAPI. All images were taken with an Olympus FV1000 confocal microscope. Antibodies used were as follows: anti-c-Fos (1:1000, guinea pig, Oasis Biofarm, Hangzhou, China), anti-GPR30 (1:500, rabbit, Alomone Labs, Isreal), anti-IBA1 (1:1000, goat, Novusbio, USA), anti-GFAP (1:1000, mouse, Cell Signaling Technology, USA), IB4-FITC (1:1000, Thermofisher, USA), Nissl (1:500, Thermofisher, USA), goat anti-guinea pig IgG-488 (1:500, Oasis Biofarm, Hangzhou, China), donkey anti-rabbit IgG-488 (1:500, Thermofisher, USA), donkey anti-rabbit IgG-555 (1:500, Thermofisher, USA), donkey anti-mouse IgG-488 (1:500, Thermofisher, USA), donkey anti-goat IgG-488 (1:500, Abcam, USA).

### *In Situ* Hybridization

To verify the specificity of transgenic mice, *In situ* hybridization was performed according to the manufacturer’s instructions from RNAscope^®^ Multiplex Fluorescent Reagent Kit v2 (Advanced Cell diagnostics, USA) with custom-designed probe for *CCK* (Mm-CCK-C1, Advanced Cell diagnostics, USA). According to the protocols, coronal lumbar spinal sections (10 μm) collected and used for fluorescence in situ hybridization to detect *CCK^+^* neurons. The slice used to stain RFP primary antibody and *CCK* probe were taken from the -80 °C refrigerator and immediately incubated with pre-cooled 4% PFA for 15 minutes, followed by gradient dehydration (50% ethanol, 70% ethanol, 100% ethanol and 100% ethanol, 5 min for each gradient). The slides were then incubated with RNAscope® hydrogen peroxide at room temperature for 10 minutes, rinsed with distilled water and PBS. Anti-RFP primary antibody (1:1000, rabbit, Rockland, USA) prepared with co-detection diluent (323180, Advanced Cell Diagnostics) was then added to the slices and incubated overnight at 4 °C for subsequent in situ hybridization staining. To characterized the specific expression of GPR30 in SDH, VGAT (Mm-VGAT-C1) and Vglut2 (Mm-Slc17a6-C2) probe were used as described above as well as anti-GPR30 primary antibody (1:250). At least 3 mice were examined and each mouse was examined at least 3 slices in these experiments.

### Real-time PCR

Mice lumbar spinal cords or DRG were collected on day 4 after CFA and on day 14 after CCI. Tissues were rapidly collected, frozen in liquid nitrogen and stored at -80℃. RNA was extracted with standard procedures using FastPure Cell/Tissue Total RNA Isolation Kit V2 (Vazyme, Nanjing, China). 500ng of total RNA from each sample was reverse-transcribed with HiScript III RT SuperMix for qPCR (+gDNA wiper) (Vazyme, Nanjing, China). Expression of each mRNA was quantified using ChamQ Universal SYBR qPCR Master Mix (Vazyme, Nanjing, China). The sequences of quantitative PCR primers were as follows: *Gper1*: F: CCTCTGCTACTCCCTCATCG, R: ACTATGTGGCCTGTCAAGGG; GAPDH: F: AAGAAGGTGGTGAAGCAGGCATC, R: CGGCATCGAAGGTGGAAGATG.

### Spinal slice preparation and whole-cell recording

Spinal slices were prepared as previously described^[57]^. Mice (6-8-week-old, 2 weeks after CCI surgery) were anesthetized with 1% pentobarbital sodium solution and perfused with ice-cold oxygenated (95% O_2_ and 5% CO_2_) cutting artificial cerebrospinal fluid (ACSF, in mM: 100 sucrose, 63 NaCl, 2.5 KCl, 1.2 NaH_2_PO_4_, 1.2 MgCl_2_, 25 glucose, and 25 NaHCO_3_), and the spinal cord was rapidly removed. Transverse spinal cord slices (300 μm, L4 to L6 segment) were prepared using a vibratome (VT1200S, Leica, Germany) and incubated in oxygenated NMDG-ACSF (in mM: 93 NMDG, 2.5 KCl, 1.2 NaH_2_PO4, 30 NaHCO_3_, 20 HEPES, 25 Glucose, 5 Na ascorbate, 2 thiourea, 3 Na pyruvate, 10 MgSO_4_ and 0.5 CaCl_2_, and adjusted to pH 7.4 with HCl) at 34 °C for 15 min. The slices were then transferred to normal ACSF (in mM: 125 NaCl, 2.5 KCl, 1.25 NaH_2_PO_4_, 1.2 MgCl_2_, 2.5 CaCl_2_, 25 glucose, and 11 NaHCO_3_) at 34 °C for 1h and maintained at room temperature before recording. The slices were transferred to a recording chamber perfused with normal ACSF saturated with 95% O_2_ and 5% CO_2_.

Whole-cell patch-clamp recordings were performed using a Heka EPC 10 amplifier (Heka Elektronik). Borosilicate glass pipettes with the resistance of 3-5 MΩ were pulled using a horizontal pipette puller (P97, Sutter instruments, USA). The pipettes were filled with cesium-based intracellular fluid (in mM: 100 CsCH_3_SO_3_, 20 KCl, 10 HEPES, 4 Mg-ATP, 0.3 Tris-GTP, 7 Tris_2_-Phosphocreatine, 3 QX-314; pH 7.3, 285–290 mOsm). Targeted whole-cell recordings were made from EGFP expressing neurons in slices taken from *CCK-Cre* mice with virus injection. For spontaneous excitatory post synaptic currents (sEPSCs) recording, the membrane potential was held at -70 mV, and for spontaneous inhibitory post synaptic currents (sIPSCs) recording, the membrane potential was held at +10 mV. Then sEPSCs and sIPSCs were recorded for 100s and analyzed using MiniAnalysis software (Synaptosoft). 5 mice per group were examined in the experiment.

To record AMPA-mediated EPSCs, an electrode placed in the deep laminae of SDH were stimulated at every 15 seconds, and the CCK^+^ neurons were voltage clamped at -70 mV. Meanwhile, the spinal slices were incubated with ACSF containing APV (100 μM) to block NMDA receptors and bicuculline (20 μM) and strychnine (0.5 μM) to block inhibitory synaptic events. AMPAR-mediated EPSCs were recorded for 15 consecutive responses after stable baseline before and after G-1 application (0.1 μM) to compare to effect of G-1 on AMPAR-mediated EPSCs. 4 mice examined in the experiment.

### Statistical analyses

All experiments were randomized. Animals were randomly chosen from multiple cages. For behavior experiments, measurements were taken blinded to condition. All data are reported as mean ± SEM. The required sample sizes were estimated on the basis of our past experience. Statistical analysis was performed using GraphPad Prism V6. Normal distribution was performed using SPSS V20. For all experiments, P<0.05 was considered to be statistically significant.

## Acknowledgements

This work was supported by the National Natural Science Foundation of China grants (82371220) and 4+X Clinical Research Project of Women’s Hospital, School of Medicine, Zhejiang University (ZDFY2022-4XA102). This work was also supported by the Fundamental Research Funds for the Central Universities (2023ZFJH01-01, 2024ZFJH01-01, and 226-2022-00227). We also thank Sanhua Fang from the Core Facilities, Zhejiang University School of Medicine for their excellent technical assistant.

## Conflict of Interest

The authors declare that they have no competing interests.

## Author contributions

X.Z.C. and Z.Z.X. conceived the study. Q.C., H.W., S.L.X., J.Q.X., L.H.X., H.L., F.F.Z., conducted the experiments and collected data. Q.C., H.W., and S.L.X. analyzed the data. Y.Y. and H.H.Z. draw the graphic abstract and revise the manuscript. A.G.D., F.X., W.X.Z., L.H.S., Q.X., L.Y.W. and C.C.J. assisted with animal maintenance and provided reagents. Q.C., X.L.Z., Z.Z.X. and X.Z.C. drafted the article. All authors approved the version to be submitted.

## Data availability

The data that support the findings of this study are available on request from the corresponding author.

## Supplemental Figure Legends

**Figure S1.**
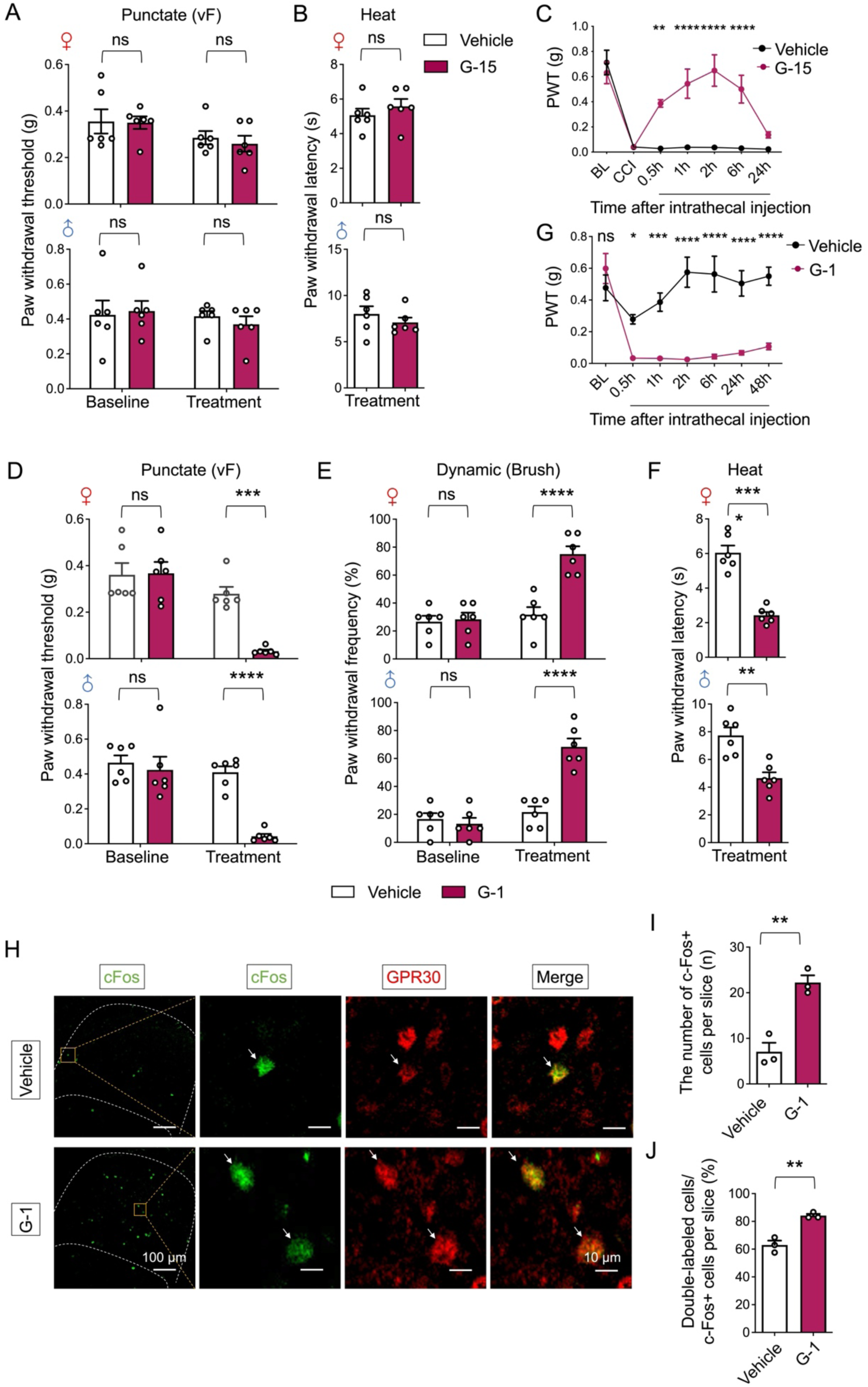
Spinal inhibition of GPR30 did not change the basic nociception while spinal activation of GPR30 mimicked neuropathic pain symptoms in naïve mice. **A, B** Behavioral tests of nociception before and after intrathecal injection of G-15 or vehicle in Von Frey tests **(A)** and Heat tests **(B)** in naive mice of both sexes (n = 6 mice for each group). **C** Time course of Von Frey tests after intrathecal injection of G-15 or vehicle in CCI mice (n = 6 mice for each group). **D-F** Behavioral tests of nociception before and after intrathecal injection of G-1 or vehicle in Von Frey tests **(D)**, Brush tests **(E)** and Heat tests **(F)** in naive mice of both sexes (n = 6 mice for each group). **G** Time course of Von Frey tests after intrathecal injection of G-1 or vehicle in naive mice (n = 6 mice for each group). **H** Immunochemical detection of c-Fos (Green) and GPR30 (Red). Scale bars: 100 μm. Boxed area of images is enlarged on the right. Scale bars: 10 μm. White arrows indicate double-positive cells. **I** Total number of c-Fos positive neurons in the SDH per section (n = 3 mice for each group, 3-5 pictures were analyzed for each mouse). **J** Percentage of c-Fos positive neurons expressing GPR30 (n = 3 mice for each group, 3-5 pictures were analyzed for each mouse). Data information: in **(A)**, ns = not significant (two-way ANOVA with Turkey’s multiple comparisons test). In **(B)**, ns = not significant (Unpaired Student’s t-test). In **(C)**, **P < 0.01; ****P < 0.0001; ns = not significant (two-way ANOVA with Turkey’s multiple comparisons test). In **(D, E)**, ***P < 0.001; ****P < 0.0001; ns = not significant. (two-way ANOVA with Turkey’s multiple comparisons test). In **(F)**, **P < 0.01; ****P < 0.0001 (Unpaired Student’s t-test). In **(G)**, *P < 0.05; ***P < 0.001; ****P < 0.0001; ns = not significant (two-way ANOVA with Turkey’s multiple comparisons test). In **(I, J)**, **P < 0.01 (Unpaired Student’s t-test). All data are presented as mean ± SEM.

**Figure S2.**
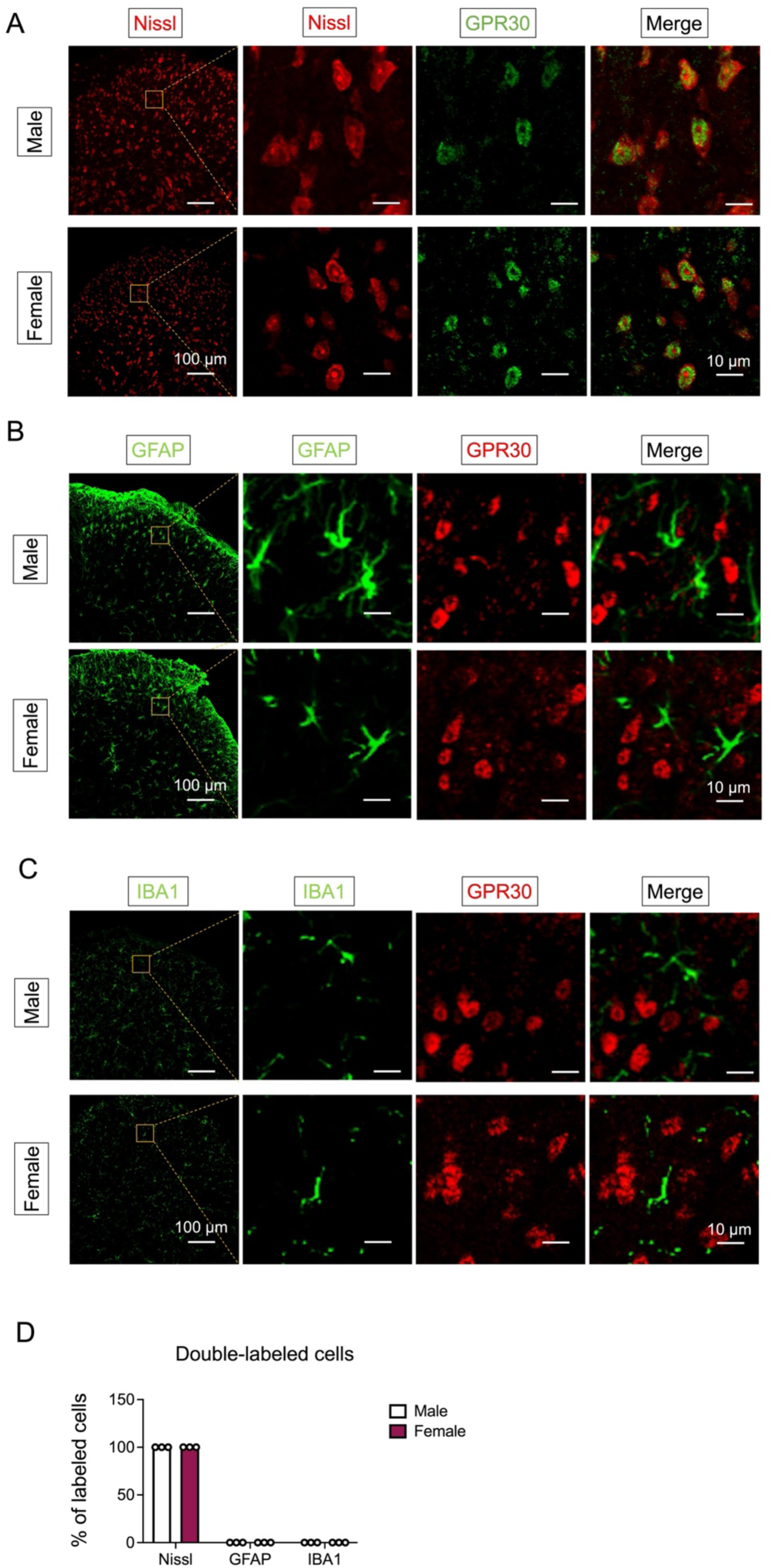
GPR30 was mainly expressed in the spinal neurons. **A** Double-staining of GPR30 (Green) with Nissl (Red) in male (up) and female mice (down). Scale bars: 100 μm. Boxed area of images is enlarged on the right. Scale bars: 10 μm. **B** Double-staining of GPR30 (Red) with GFAP (Green) in male (up) and female mice (down). Scale bars: 100 μm. Boxed area of images is enlarged on the right. Scale bars 10 μm. **C** Double-staining of GPR30 (Red) with IBA1 (Green) in male (up) and female mice (down). Scale bars: 100 μm. Boxed area of images is enlarged on the right. Scale bars: 10 μm. **D** Percentage of GPR30 expressed in neurons, astrocytes and microglia in both sexes of mice (n = 3 mice for each group, 3 pictures were analyzed for each mouse). Data information: in **(D)**, data are presented as mean ± SEM.

**Figure S3.**
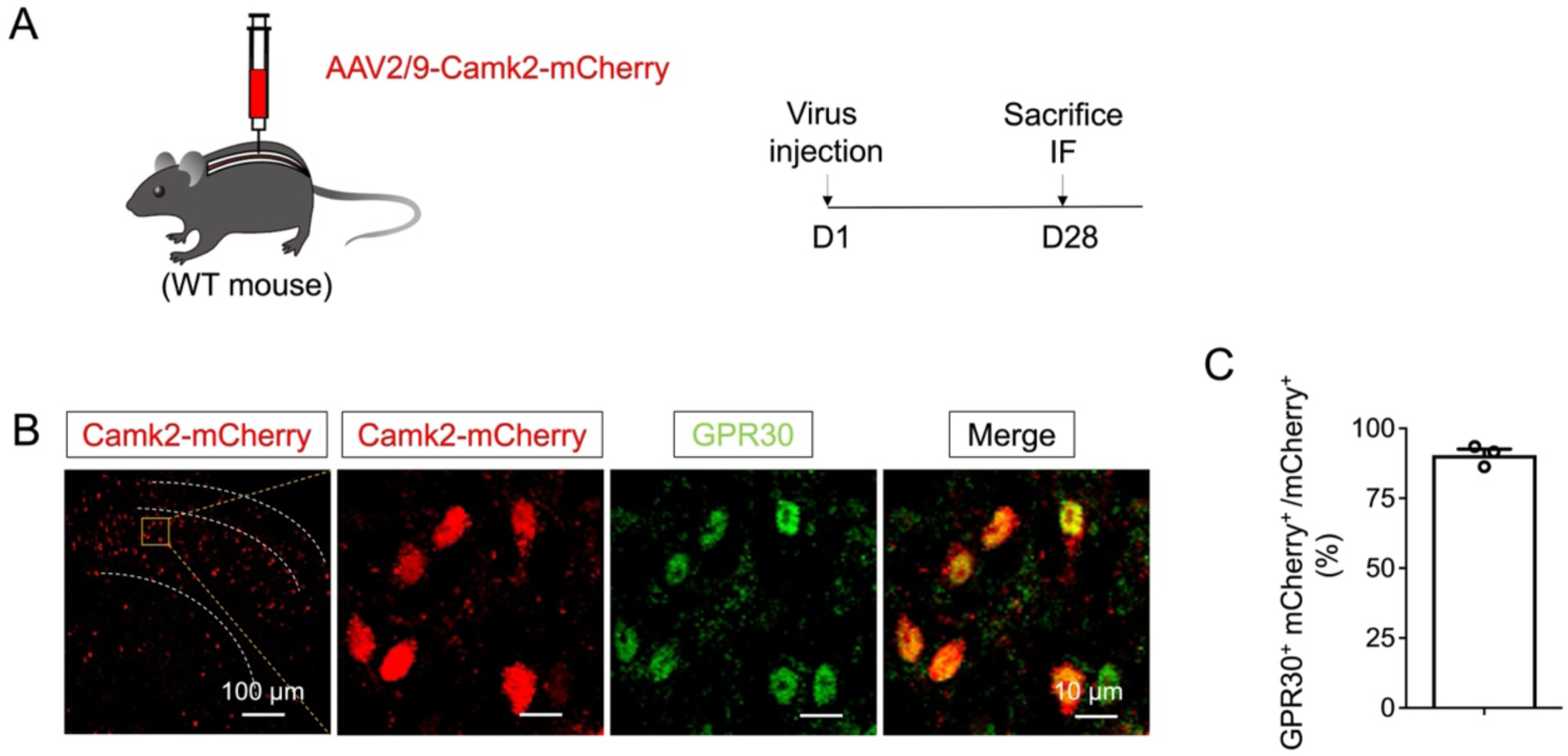
GPR30 was widely expressed in the Camk2^+^ excitatory interneurons in the SDH. **A** Schematic illustration of the strategy to identify the Camk2^+^ excitatory interneurons in spinal dorsal horn by AAV. **B** Double staining of Camk2^+^ mCherry^+^ neurons (Red) with GPR30 (Green) by immunohistochemistry. Scale bars: 100 μm. Boxed area of images is enlarged on the right. Scale bars: 10 μm. **C** Percentage of Camk2 positive neurons expressing GPR30 in wild mice. (n = 3 mice, 3-4 pictures were analyzed for each mouse)

**Figure S4.**
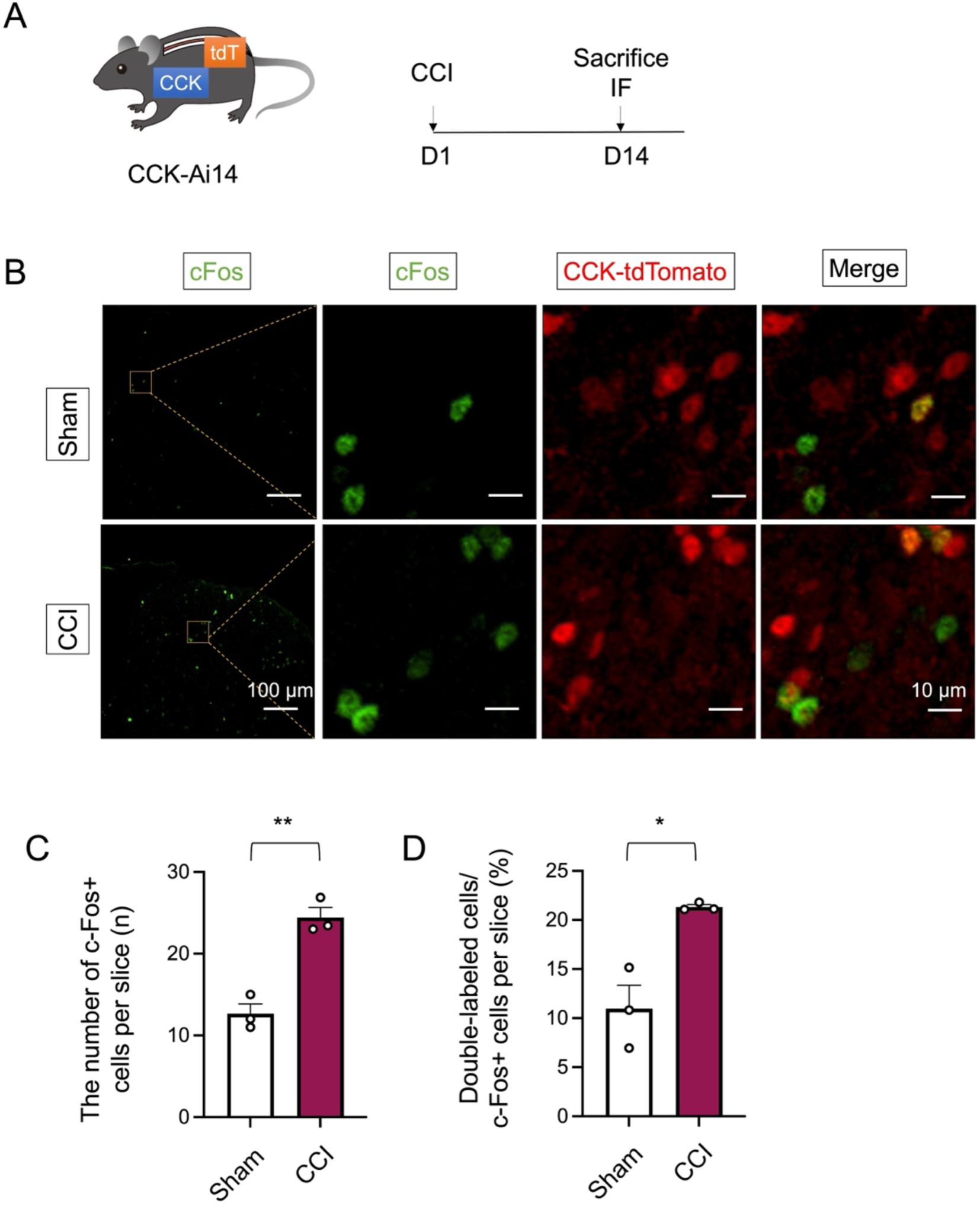
CCK positive neurons were more activated after CCI. **A** Schematic illustration of CCK-Ai14 subjected with CCI. **B** Immunochemical detection of c-Fos (Green) and CCK-tdTomato (Red). Scale bars: 100 μm. Boxed area of images is enlarged on the right. Scale bars: 10 μm. **C** Total number of c-Fos positive neurons in the SDH per section (n = 3 mice for each group, 6-8 pictures were analyzed for each mouse). **D** Percentage of c-Fos positive neurons expressed on CCK-tdTomato positive neurons (n = 3 mice for each group, 6-10 pictures were analyzed for each mouse).

**Figure S5.**
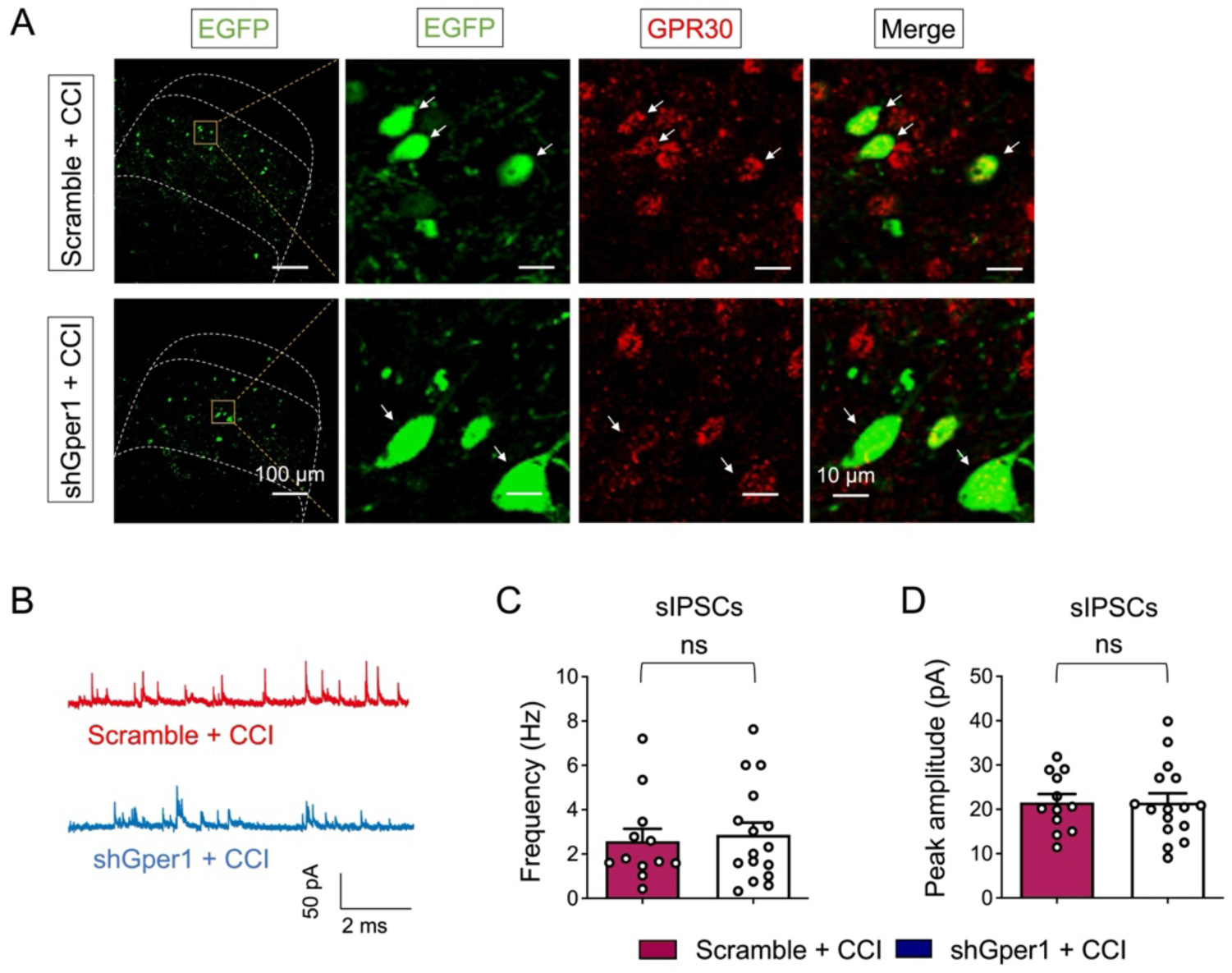
Knockdown of GPR30 in spinal CCK^+^ neurons did not change the sIPSC in CCI mice. **A** Immunohistochemical detection of GPR30 (Red) and shRNA (Green) in mice with virus injection. Note that the intensity of GPR30 fluorescence is less in shGper1-EGFP^+^ cells than that in scramble-EGFP^+^ cells. Scale bars: 100 μm. Boxed area of image is enlarged on the right. Scale bars: 10 μm. **B** Representative spontaneous IPSCs in CCK-EGFP neurons. **C** The frequency of spontaneous IPSCs (n = 12-16 cells from 5 mice). **D** The peak amplitude spontaneous IPSCs (n = 12-16 cells from 5 mice). Data information: in **(C, D)**, ns = not significant (Unpaired Student’s t-test). All data are presented as mean ± SEM.

**Figure S6.**
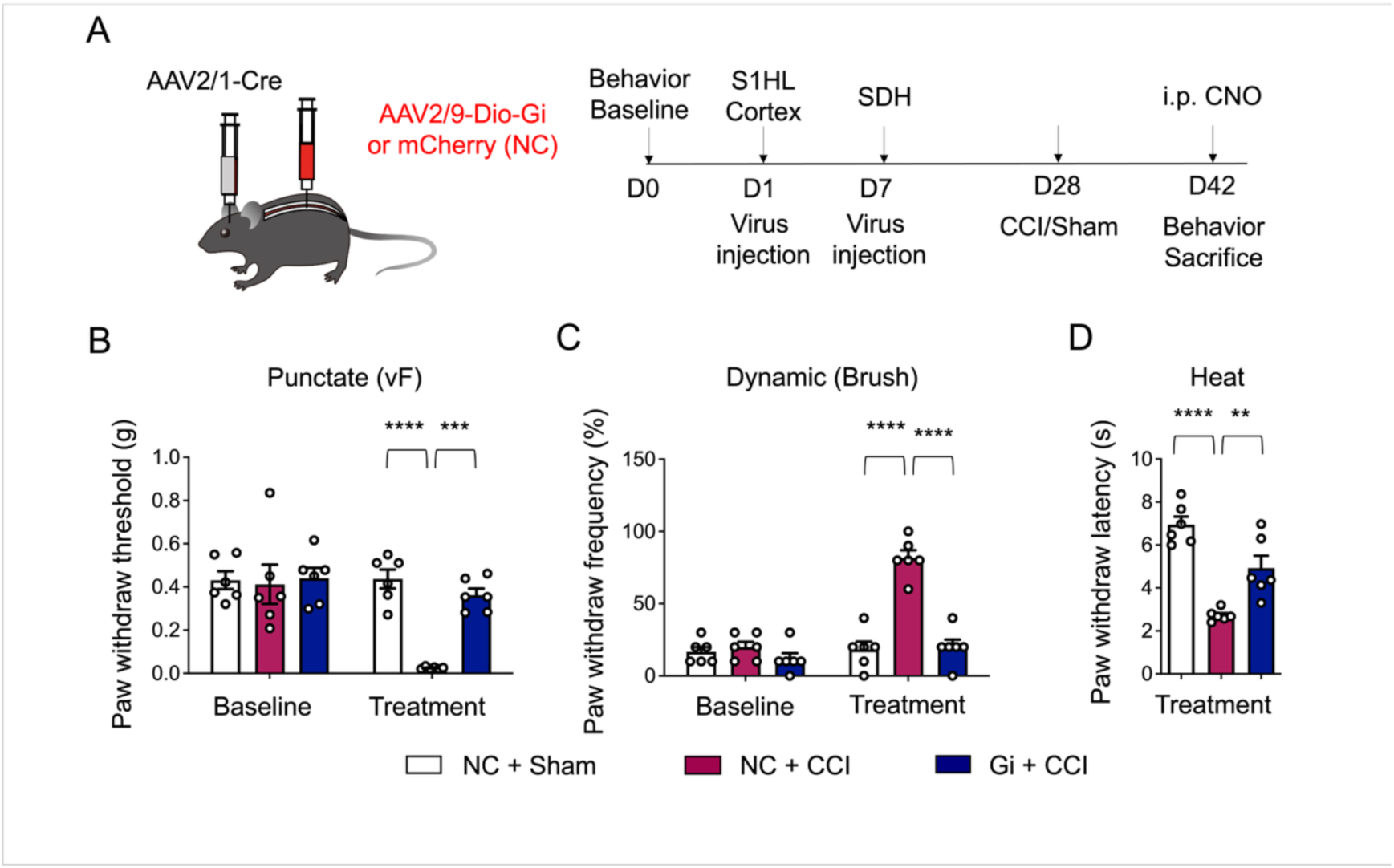
CCI-induced neuropathic pain was relieved by chemogenetic inhibition of S1-SDH post-synaptic neurons. **A** Schematic illustration of the strategy for identifying the post-synaptic neurons of S1HL-SDH projections in SDH (Left) and diagram showing the timeline of AAV2/1 injection in the S1HL cortex, AAV2/9 injection in the spinal dorsal horn, CCI surgery and behavioral tests (Right). **B-D** Behavioral tests of basic nociception, 14 days after CCI surgery along with intraperitoneal injection of CNO in Von Frey tests **(B)**, Brush tests **(C)** and Heat tests **(D)** in mice (n = 6 mice per group). Data information: In **(B, C)**, ***P < 0.001; ****P < 0.0001. (two-way ANOVA with Turkey’s multiple comparisons test). In **(D)**, **P < 0.01; ****P < 0.0001 (one-way ANOVA with Turkey’s multiple comparisons test). All data are presented as mean ± SEM

## Supplemental Movie Legends

**Movie 1. Heat test of mice subjected with CCI and spinal viral injection.**

**Movie 2. VonFrey test of mice subjected with CCI and spinal viral injection.**

**Movie 3. Pre-stimulation phase of RT-PEA test of mice subjected with CCI and shGper1.**

**Movie 4. Stimulation phase of RT-PEA test of mice subjected with CCI and shGper1.**

**Movie 5. Post-stimulation phase of RT-PEA test of mice subjected with CCI and shGper1.**

**Movie 6. Pre-stimulation phase of RT-PEA test of mice subjected with CCI and Scramble.**

**Movie 7. Stimulation phase of RT-PEA test of mice subjected with CCI and Scramble.**

**Movie 8. Post-stimulation phase of RT-PEA test of mice subjected with CCI and Scramble.**

## Notes

### Competing Interest Statement

The authors have declared no competing interest.

### Summary of Updates

we have revised the manuscript for improved clarity according to the latest comments from reviewers.

